# Dissociable norepinephrine and dopamine representations during strategic decision-making and reward learning

**DOI:** 10.64898/2026.10.31.763226

**Authors:** Joanna Aloor, Timothy PH Sit, Joschua Geuter, Joseph O Tutt, Chunyu A Duan

**Affiliations:** Sainsbury Wellcome Centre for Neural Circuits and Behaviour, University College London, London, UK

**Keywords:** multi-player game, strategic decision-making, norepinephrine, dopamine, stochastic choice, learning, pupil

## Abstract

Adaptive decision-making in competitive environments requires learning from immediate reward outcomes while tracking longer-timescale information to adjust behavioural strategies and remain unpredictable against others. Dopamine (DA) and norepinephrine (NE) have long been implicated in reward-guided behaviours, but their contributions to strategic choice remain unclear. We trained mice to play a zero-sum matching pennies game against a computer opponent that exploited regularities in their choice and reward history, such that optimal performance required stochastic choice decoupled from immediate reward. We combined fibre photometry recordings, behavioural and neural modelling, and a within-subject multi-task design to determine how NE and DA contribute to strategic choice in the matching pennies game versus reward-guided behaviour in a two-armed bandit (2AB) task. DA responses were dominated by trial-by-trial reward outcomes and consistently tracked model-derived reward prediction errors (RPE) across tasks and behavioural states. In contrast, NE activity reflected salient task events, predicted upcoming choice switches, and tracked longer-timescale variables including model-inferred behavioural strategy and average reward rate. In addition, NE responses were selectively enhanced following rewarded switches when these outcomes were informative for deciding whether to stay in or switch from the current strategy; and this enhancement could not be explained by RPE. Incorporating pre-choice NE activity into a reinforcement-learning model to modulate choice generation improved model prediction in matching pennies but not in the 2AB task. Together, these findings reveal complementary neuromodulatory correlates of competitive decision-making across timescales: DA primarily guides learning from immediate outcomes through RPE, whereas NE additionally represents contextual information and choice stochasticity relevant for shaping long-term strategies.

## Introduction

To maximise reward in competitive environments, animals are not only guided by reward learning, but also need to integrate longer timescale information and form strategies that may be decoupled from immediate action-outcome associations. For example, while it is important to learn the location of rich food sources during competitive foraging and to know the safest path to shelter when escaping from predators, it is also essential to behave stochastically so as not to be easily predicted by others. Game-theoretic tasks provide a tractable framework for studying this form of strategic decision-making by requiring individuals to select actions and simultaneously consider and adapt to how these actions may be used against them (Camerer, 2003; Nash, 1950; von Neumann and Morgenstern, 1944). Specifically, competitive mixed-strategy games such as matching pennies (Barraclough, Conroy, and Lee, 2004) create such a computational demand that is often not exposed in conventional reward-guided decision-making tasks: optimal behaviour requires regulating choice stochasticity itself, independently of the value assigned to individual actions (Barraclough, Conroy, and Lee, 2004; Lee et al., 2004; Tervo et al., 2014; Wang et al., 2022).

This distinction separates strategic decision-making from the reinforcement-learning framework that has dominated the study of reward-guided behaviour. In standard reinforcement learning, an agent learns a mapping between actions and their expected consequences and uses prediction errors to update these value estimates (Sutton and Barto, 2018). Dopamine (DA) has provided one of the clearest neural implementations of this computational framework. Midbrain dopaminergic neurons respond to reward prediction errors (RPEs), signalling discrepancies between obtained and predicted outcomes in a manner consistent with temporal-difference learning (Bayer and Glimcher, 2005; Cohen et al., 2012; D’Ardenne et al., 2008; Montague, Dayan, and Sejnowski, 1996; Schultz, Dayan, and Montague, 1997). Dopamine release in the nucleus accumbens (NAc) exhibits closely related relationships to reward prediction and outcome, supporting a conserved role for mesolimbic DA in reinforcement learning (Day et al., 2007; Hart et al., 2014). More recent work has expanded the RPE framework to account for richer forms of DA signalling, including distributional value coding (Dabney et al., 2020), ‘feature-specific RPE’ (Lee et al., 2024), prediction errors for perceived causal control over learned associations (Bernklau et al., 2024), and action prediction errors supporting value-free learning (Greenstreet et al., 2025). These findings have established a detailed account of how DA signals can support updating of value representations from experience.

In competitive environments, RPE signals alone are insufficient to guide optimal behaviour. When an opponent exploits predictable patterns in an animal’s choices, optimal performance requires the animal to increase or decrease the variability of its choices irrespective of the expected value of the available actions. This type of stochasticity is commonly formalised in reinforcement-learning models by the inverse temperature of a softmax choice rule, which determines the extent to which an animal’s estimated values determine its choice (Daw, Niv, and Dayan, 2005; Sutton and Barto, 2018). Yet compared with the extensive circuit-level literature describing how neural signals update action values, the neural mechanisms that regulate choice stochasticity on a trial-by-trial basis remain poorly understood.

Norepinephrine (NE), released throughout the forebrain by locus coeruleus (LC) neurons, has long been proposed to regulate how behaviour adapts to changing environmental demands. Meta-learning accounts propose that neuromodulatory systems adjust learning rates according to environmental stability (Doya, 2002), while adaptive gain and Bayesian theories propose that LC-NE regulates exploration-exploitation by signalling changes in uncertainty and task contingencies (Aston-Jones and Cohen, 2005a; Aston-Jones and Cohen, 2005b; Dayan and Yu, 2006; O’Reilly et al., 2013; Yu and Dayan, 2005). Related network-reset models suggest that phasic LC activity can reconfigure ongoing processing when salient events require a change in behavioural state (Bouret and Sara, 2005; Sara and Bouret, 2012). These frameworks suggest that NE may regulate the flexibility with which behavioural policies are engaged and updated in response to changing information.

A growing experimental literature provides support for aspects of these predictions. Causal manipulations of NE signalling have implicated the LC-NE system in uncertainty, exploration, behavioural variability and flexibility (Breton-Provencher et al., 2022; McBurney-Lin et al., 2022; Tervo et al., 2014; Wang et al., 2025). While these studies are consistent with a role of NE in regulating strategic stochasticity, LC-NE activity itself has not been directly mapped onto trial-by-trial behavioural strategy to probe the mechanism through which NE facilitates competitive decision-making. At the same time, recent work indicates that LC-NE signalling can contain prediction-error-like information, challenging a simple division in which DA represents RPE while NE represents uncertainty or arousal (Su et al., 2026). LC-NE neurons have also been reported to show functional heterogeneity and organised topographically, with subsets of dorsal LC neurons responding to choice switches and RPEs, while other LC populations show different relationships to reward (Su et al., 2026). Comparisons of NE and DA during strategic choice versus reward-guided learning can provide new insights on their distinct contributions to adaptive behaviours across timescales.

Here, we investigated DA and NE dynamics in a mouse matching pennies task where a computer opponent exploited regularities in the animals’ choice and reward history, and the optimal strategy was to choose between two options stochastically. Mice initially employed structured choice biases but progressively adopted a more stochastic strategy. We recorded DA release in the nucleus accumbens and LC-NE activity whilst animals learned the task. We found that DA responses were dominated by trial-by-trial outcome signals and were consistently accounted for by RPEs derived from a reinforcement learning (RL) model. In contrast, NE activity reflected salient task events, predicted choice switching, and encoded longer-timescale information such as model-inferred behavioural strategy and average reward rate. Furthermore, NE responses to rewarded choice switches were enhanced only when these outcomes were informative for deciding whether to stay in or switch from the current strategy, and this effect could not be accounted for by RPEs alone. Finally, scaling the stochasticity of choice policy in our RL model with trial-by-trial NE activity significantly improved model performance. Together, these findings identify complementary contributions of DA and NE to competitive decision-making, with DA guiding reward learning through RPEs and NE representing broader contextual and strategic variables that shape flexible deployment of behaviours in complex decision scenarios.

## Results

### Norepinephrine and dopamine primarily encode salient task events and immediate rewards respectively

To test the roles of norepinephrine (NE) and dopamine (DA) in strategic decision-making, we trained head-fixed mice on a two-choice competitive task called matching pennies (Figure 1a; Aloor et al., 2026), whilst measuring NE and DA dynamics using fibre photometry. In the matching pennies task, mice played against a computer opponent that aimed to predict their upcoming choice based on statistical regularities in their choice and reward history. Each trial began with a sound cue, followed by extension of motorised lick ports that remained available during the choice and reward period (Figure 1a). Mice could choose either the left or right port and received a reward immediately if their choice was not predicted by the computer opponent. Reward was omitted if the computer prediction was correct. Across weeks of training, mice transitioned from highly structured behaviour, predominantly favouring one side, to more variable and unpredictable choices that approached the optimal 50% reward rate (Figure 1b-d, Figure S1a-c). We characterised these changes in behaviour using a hidden Markov model (HMM), in which latent states captured distinct decision-making strategies defined by different combinations of generalised linear model (GLM) inputs (Figure 1e; Aloor et al., 2026; Ashwood et al., 2022; Bolkan et al., 2022). Using three GLM inputs (side bias, repetition of the previous rewarded choice, and repetition of the previous unrewarded choice), we found that three states were sufficient to describe mouse behaviour (Figure S1d). These states reflected two biased strategies, favouring the right and left sides, and a stochastic strategy that used more variable and unpredictable choices (Figure 1f, Figure S1e). Across learning, mice progressively increased the use of the stochastic strategy (Figure 1g, Figure S1f), providing a behavioural framework for investigating the neuromodulatory mechanisms that control adaptive stochasticity while mice improved their strategic gameplay.

**Figure 1.**
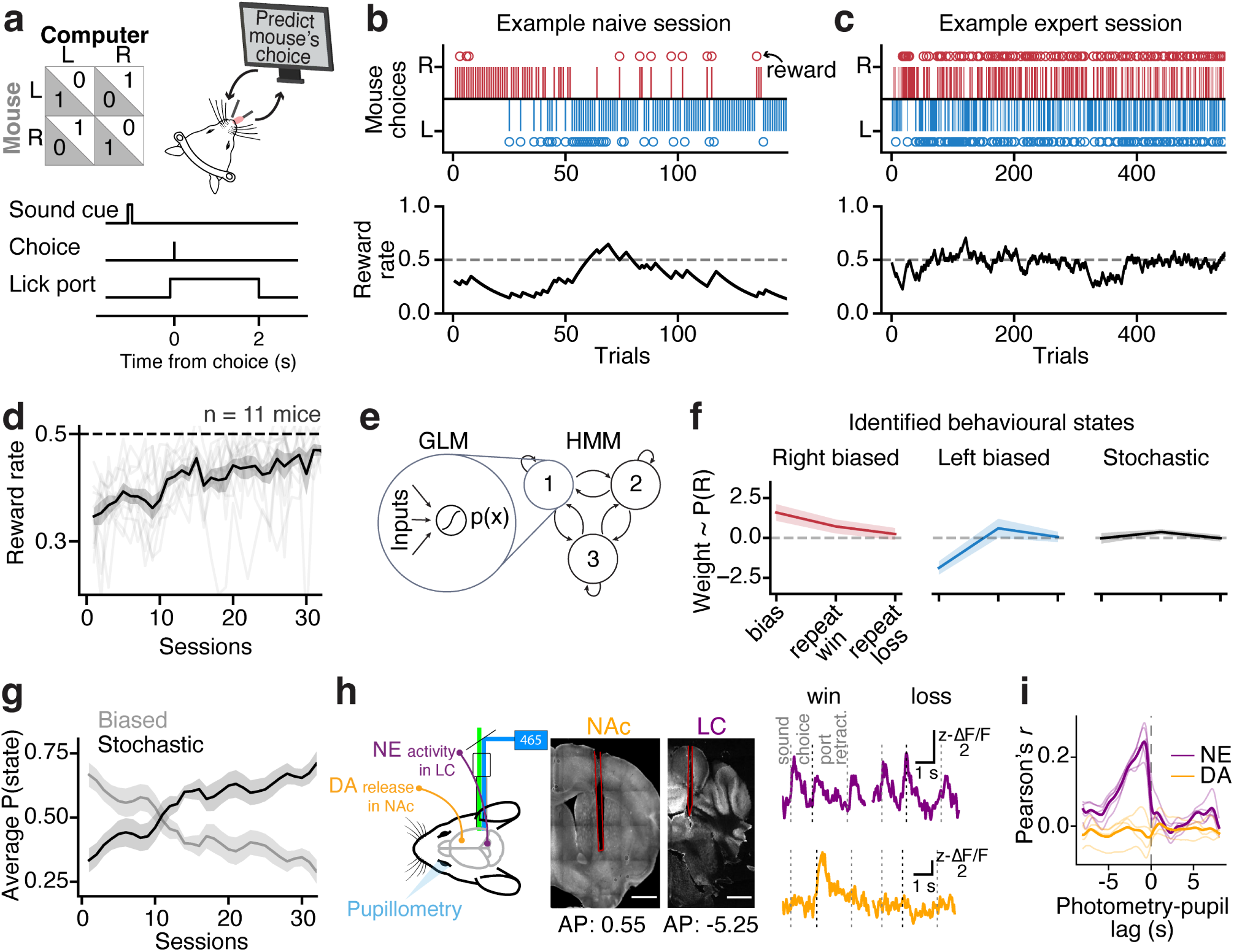
Fibre photometry recordings of norepinephrine and dopamine whilst mice learned a multi-player competitive decision-making task. **a)** Matching pennies task schematic and trial design. **b)** Example naive session early in learning showing trial-by-trial choices (top) and smoothed reward rate (bottom). **c)** Similar to (b), showing performance of the same animal during an example expert session. **d)** Reward rate across learning. Thick line indicates mean *±* SEM across animals; thin lines represent individual mice (n=11 mice). **e)** GLM-HMM used to identify behavioural strategy states across learning. **f)** GLM weights for a three-state GLM-HMM model fit to mouse behaviour. Lines represent mean *±* SEM across model fits for individual animals. GLM weights were grouped across mice using constrained K-means. **g)** Mean P(state) across learning for the stochastic and biased (left and right biased combined) states. Lines represent mean *±* SEM across animals. **h)** Fibre photometry recordings of GCaMP8s in locus coeruleus (LC) norepinephrine (NE) neurons and dLight capturing dopamine (DA) release in nucleus accumbens (NAc). Left, schematic for experimental setup. Right, example histology showing fibre placement in NAc (left) and LC (right). Fibre location is outlined in red. Scale bar = 1 mm. **i)** Cross-correlation of pupil with NE/DA. Thick line represents mean across animals; thin lines indicate individual mice.

We recorded pupil responses, LC-NE activity in Dbh-Cre mice expressing GCaMP8s in the locus coeruleus (n = 5 mice), and DA release in the nucleus accumbens using the genetically encoded dopamine sensor dLight1.1 (n = 6 mice; Figure 1h, Figure S1g). Previous studies have shown that pupil dynamics closely track norepinephrine activity, raising the possibility that pupil diameter could provide a non-invasive readout of NE-linked changes in arousal and other cognitive states (Gilzenrat et al., 2010; Joshi et al., 2016; Reimer et al., 2016). Consistent with these reports, we found that changes in pupil size were strongly correlated with NE activity, with the pupil response lagging NE by approximately 1–2 s, whereas DA activity showed much weaker relationships with pupil dynamics (Figure 1i). Additionally, both NE and pupil responses after the outcome period were positively correlated with response time (RT) on the next trial; whereas post-outcome dopamine activity was negatively correlated with RT on the next trial (Figure S2a-d), even when controlling for other variables that influenced RTs (Figure S2e-h).

Given the established role of dopamine and norepinephrine in encoding reward information (Dabney et al., 2020; Su et al., 2026; Watabe-Uchida, Eshel, and Uchida, 2017), we first asked how these neuromodulators differentially represent immediate reward outcomes. As expected, DA responses showed a strong separation between wins and losses at the time of reward delivery (Figure 2a). While NE activity also differentiated reward outcomes, although less strongly than DA, its activity was dominated by responses to key events occurring throughout the trial (Figure 2b). Three prominent peaks in NE activity aligned with the trial-start sound cue, the animal’s choice, and retraction of the lick ports following the outcome, suggesting that NE is more broadly engaged by salient task events rather than selectively encoding reward outcome. These patterns of responses were consistent across the biased and stochastic states identified by GLM-HMM (Figure S3a-d).

**Figure 2.**
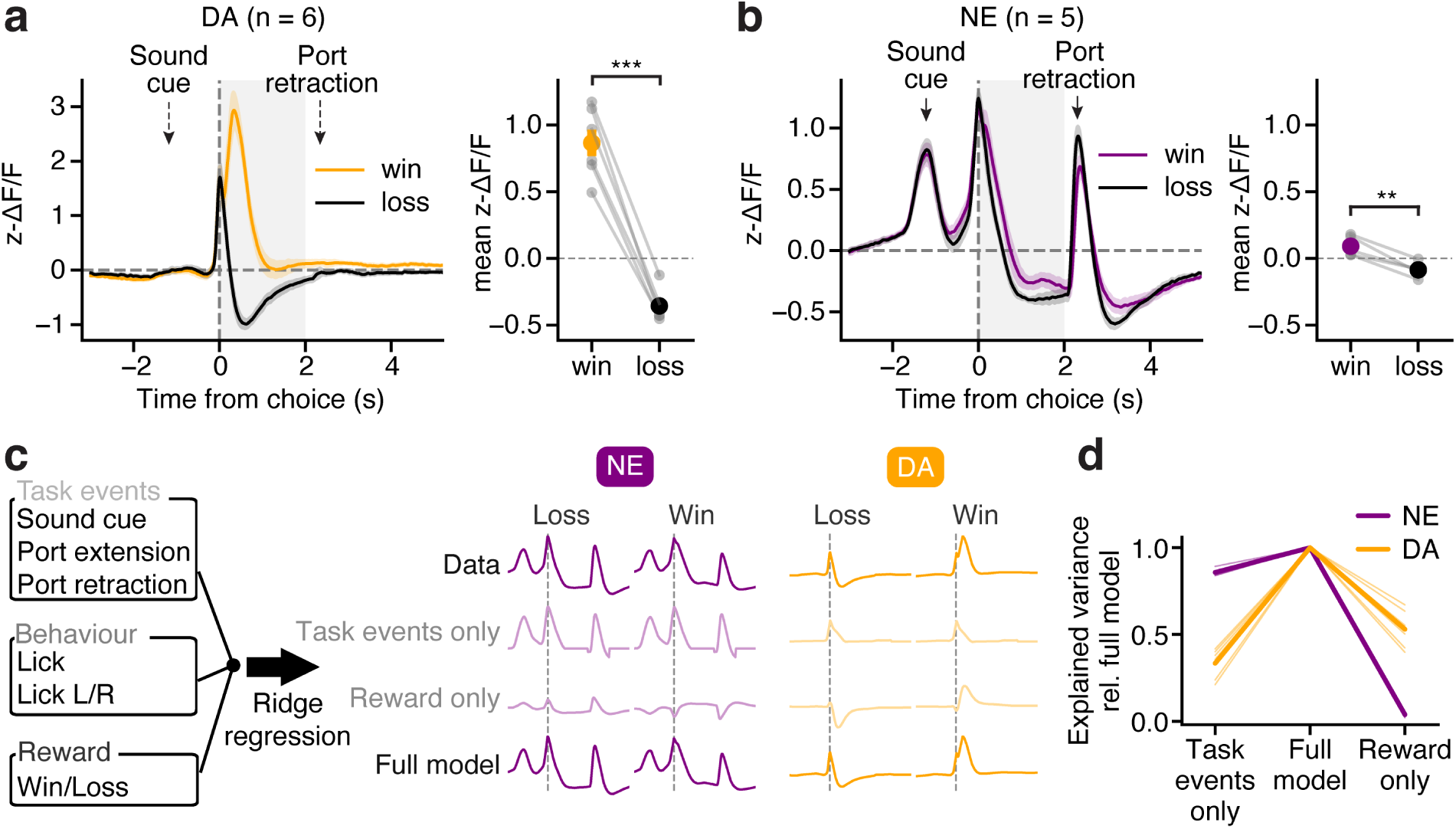
Dopamine and norepinephrine encoding of immediate reward outcomes and salient task events. **a)** DA response to current trial reward outcome (left) and difference between the averaged post-choice (0 – 2 s) DA response in win and loss conditions. Trials used from all sessions across learning. Paired t-test across animals, *p* = 3.37*×* 10*^−^*^5^, n = 6 mice; win mean = 0.865, loss mean = -0.356. Lines indicate mean *±* SEM across animals (n=6 DA mice, n=5 NE mice). **b)** Same as (a) for NE activity. Paired t-test across animals, *p* = 0.0027, n = 5 mice; win mean = 0.091, loss mean = -0.086. **c)** Linear encoding model predicting NE/DA responses using regressors for task events, behaviour, and reward. Dashed line indicates time of choice. **d)** Difference in explained variance of a partial model containing only task events regressors or only reward regressors relative to the full model described in (c). Thin lines show individual mice and thick lines show mean across mice (N = 5 mice for NE, N = 6 mice for DA). ** p<0.01, *** p<0.001

To quantify these differences while controlling for other task-relevant features, we fit linear encoding models predicting NE and DA activity from regressors representing task events (sound cue and port movement), behaviour (lateralised licking), and reward outcome (win/loss) (Figure 2c). We compared the full model with two partial models containing either task-event or reward regressors alone. This analysis confirmed that NE activity was explained predominantly by salient task events, with relatively little additional contribution from reward outcome, whereas DA activity responded more to reward-related information (Figure 2d), with some degree of explained variance from task events that is likely attributed to the close timing between port extension and reward consumption. Since NE and pupil signals are correlated, we further tested whether NE carried unique information beyond pupil-related signals. Comparing models with time-lagged pupil regressors plus task variables revealed that NE still contained significant task-related information unexplained by pupil (Figure S2i-l). In contrast, pupil explained little variance in DA signals, consistent with their low correlation (Figure 1i).

### Pre-choice norepinephrine, but not dopamine, reflects longer-timescale reward and strategy information

In the matching pennies game, maximising long-term reward requires animals to avoid over-weighting the outcome of any single trial and instead integrate information over trials. Given that DA strongly encodes immediate outcome but NE shows relatively little outcome-related modulation, we next asked whether NE signals instead track reward and strategy information over longer timescales. Specifically, we tested the relationship between neuromodulatory dynamics and average reward rate (Methods), GLM-HMM-inferred behavioural strategies, and the probability of upcoming choice switching (Figure 3a). An exponentially weighted moving average was used to compute a locally smoothed estimate of average reward rate per trial. We then divided average reward rate into five equal-sized quantiles across sessions and characterised NE and DA responses to trials within each quantile. NE activity varied systematically with average reward rate, with a strong positive relationship during the time preceding choice (Figure 3b). By averaging activity within three epochs spanning pre-choice (-2 – 0 s), post-choice (0 – 2 s), and post-outcome (2 – 4 s) periods, we demonstrated that NE activity was positively correlated with average reward rate before choice, but negatively correlated with average reward rate following choice and outcome (Figure 3c).

**Figure 3.**
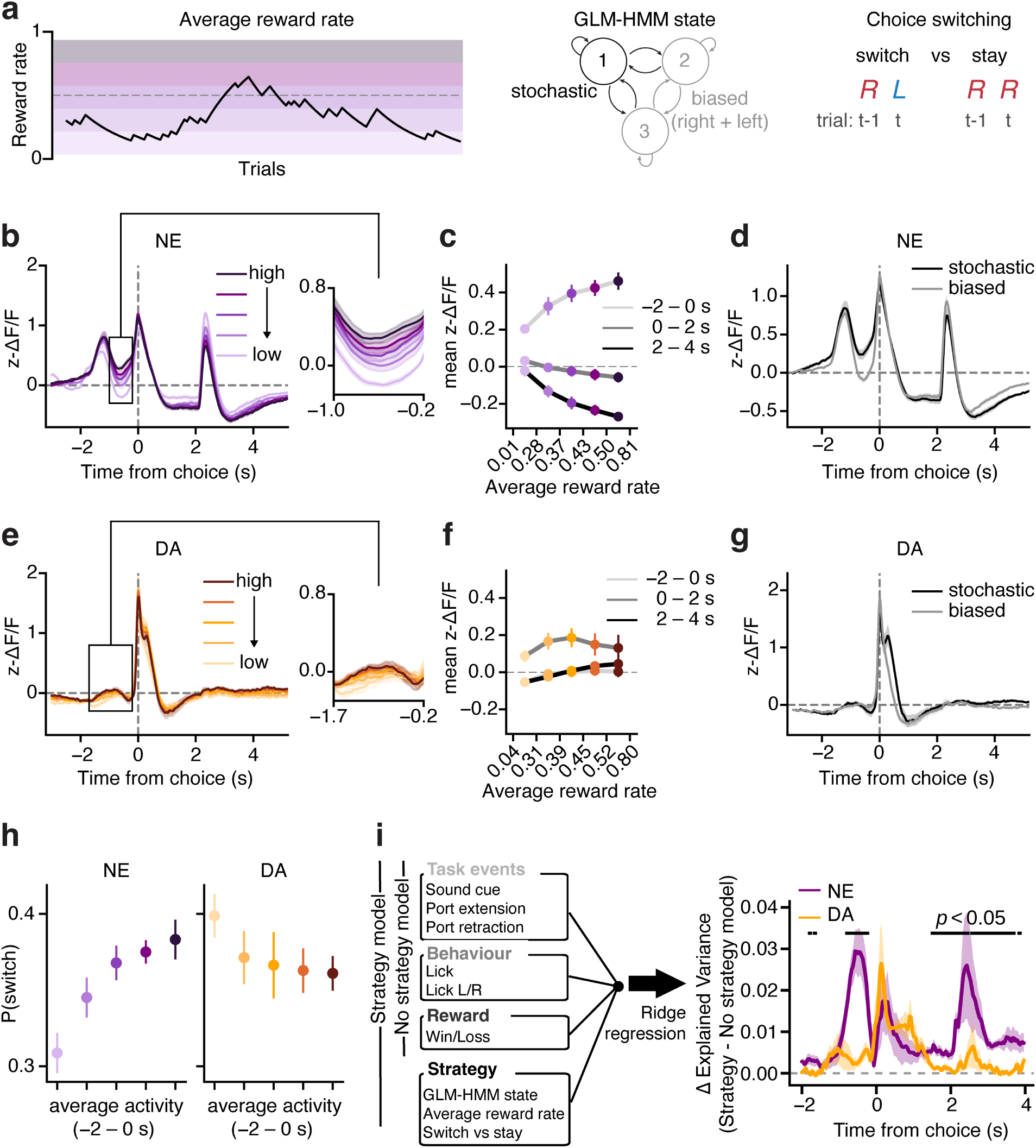
Norepinephrine tracked longer timescale reward and strategy information. **a)** We tested if NE and DA encode three different examples of strategy-related information: average reward rate calculated using an exponential smoothing kernel with a time constant, *τ ≈* 20 trials (left), GLM-HMM state (middle), and choice switching (right). **b)** Average NE activity of trials split into five quantiles based on smoothed reward rate. Black box indicates the time period of the zoomed-in pre-choice activity (left). Lines show mean *±* SEM across mice (n=5 NE mice, n=6 DA mice). **c)** Average NE activity across three trial epochs for each smoothed reward rate quantile. Data points indicate mean *±* SEM error bars across mice (n=5 NE mice, n=6 DA mice). Pre-choice: one-sample t-test of per-subject OLS slopes (activity *∼* reward rate) against 0, *p* = 0.0011, n = 5 mice; mean slope = 0.689. Post-choice: *p* = 0.131; mean slope = -0.244. Post-outcome: *p* = 0.0037; mean slope = -0.678. **d)** NE response separating trials belonging to the stochastic (black) and biased (grey) states. Lines show mean *±* SEM across mice. **e-g)** Same as (b-d) for DA responses. Pre-choice: one-sample t-test of per-subject OLS slopes (activity *∼* reward rate) against 0, *p* = 0.0262, n = 6 mice; mean slope = 0.187. Post-choice: *p* = 0.493; mean slope = 0.122. Post-outcome: *p* = 0.0287; mean slope = 0.300. **h)** P(switch) of the upcoming choice predicted by averaged pre-choice activity (-2 – 0 s) split into five quantiles. Data points indicate mean *±* SEM error bars across mice. **i)** Difference in explained variance of the linear encoding model when adding strategy regressors (GLM-HMM state, average reward rate, and switching). Lines show mean and shading shows *±* 1 SEM across mice (N = 5 mice for NE, N = 6 mice for DA). Black bars indicate time bins where there is a significant difference between DA and NE (two-sided Wilcoxon rank-sum test per bin, p < 0.05).

Pre-choice NE responses also differed according to the animal’s behavioural strategy, with higher activity in the stochastic state compared to the biased state (Figure 3d). This strategy-related information in NE was present when controlling for preparatory movement (Figure S4) as well as differential state occupancy in early versus late learning stages (Figure S5a-c). Furthermore, we tested whether within-session task engagement could underlie the apparent state encoding (Figure S5g-l). We found that behavioural changes as a function of session progression were consistent with increased satiation and decreased engagement towards the end of a session. However, the biased state and not the stochastic state was more correlated with disengagement. We then matched the number of trials in the stochastic state and biased state for each position bin within a session (Figure S5k), and still found significant GLM-HMM state selectivity in pre-choice NE activity (Figure S5l). In contrast, DA activity showed weaker and less systematic relationship with average reward rate or GLM-HMM-inferred state across all three trial epochs (Figure 3e-g), with or without controlling for movement, learning, and task engagement (Figure S4, Figure S5).

Because behavioural state and reward rate are themselves correlated with each other, we also examined neuromodulatory responses to average reward rate while controlling for GLM-HMM state and found consistent results (Figure S3e-h). A linear regression model predicting NE responses from reward rate, state, and their interaction confirmed that pre-choice NE activity was significantly explained by both reward rate (mean *β* = 0.29 *±* 0.07 SEM, *p* = 0.034) and behavioural state (*β* = 0.08 *±* 0.02, *p* = 0.034), with no significant interaction (*p* = 0.797; Figure S3g). These results indicate that reward rate and behavioural state contributed additively to pre-choice NE activity. The same regression applied to pre-choice DA activity found no significant effect of reward rate (*p* = 0.207), state (*p* = 0.068), or their interaction (*p* = 0.207).

To avoid predictable patterns that can be exploited by the opponent, it is important to vary choices over successive trials. We therefore asked whether pre-choice neuromodulatory activity could predict choice switching on that trial. We divided trials into five quantiles according to pre-choice activity (-2–0 s) and calculated the probability of switching of the upcoming choice. NE activity was positively associated with choice switching, such that higher pre-choice NE activity predicted greater probability of switching, whereas DA activity showed the opposite relationship (Figure 3h). To test whether this relationship held when behavioural state was taken into account, we fit a logistic regression model predicting switching from pre-choice activity, behavioural state, and previous trial outcome, separately for each animal. Both pre-choice NE activity (mean *β* = 0.10 *±* 0.04 SEM, one-sample t-test, *p* = 0.045) and behavioural state (*β* = 1.67 *±* 0.23, *p* = 0.002), but not previous trial outcome (*p* = 0.307), were significantly correlated with switching, suggesting that NE activity predicts choice switching independently of behavioural state and recent reward history.

Finally, we confirmed these strategy-related variables significantly improved explained variance of NE signals beyond the task events and immediate reward outcomes by extending our linear encoding model to include GLM-HMM state, average reward rate, and upcoming choice switch as additional regressors. This result was observed for NE during both the pre-choice and post-outcome periods, whereas the corresponding improvement was substantially weaker for DA (Figure 3i). Together, these findings indicate that NE carried information about longer-timescale reward statistics and tracked model-inferred behavioural strategy, whereas these signals were largely absent from DA activity, which remained strongly anchored to the encoding of immediate reward outcomes.

### Post-choice norepinephrine enhances choice–outcome information when relevant for strategy updating

Norepinephrine has previously been proposed to promote behavioural switching when changes in the environment require an animal to update its current policy (Bouret and Sara, 2005; Tervo et al., 2014). Our behavioural framework provides an opportunity to test this hypothesis by comparing the neural representation of choice and reward under distinct strategies in which reward outcomes provide inherently different amounts of information for strategy evaluation. We compared trials in which mice either switched or repeated their previous choice (stay), separately for rewarded and unrewarded outcomes and for biased and stochastic states (Figure 4a). Importantly, the informativeness of these choices and outcomes differs between states (Figure 4b). In the stochastic state, rewards are distributed approximately randomly between left and right choices, and receiving a reward after either switching or staying should provide little information about which choice should be favoured on subsequent trials. In contrast, during the biased state, receiving a reward following a switch could provide evidence that the newly selected choice is more advantageous, at least in the short term.

**Figure 4.**
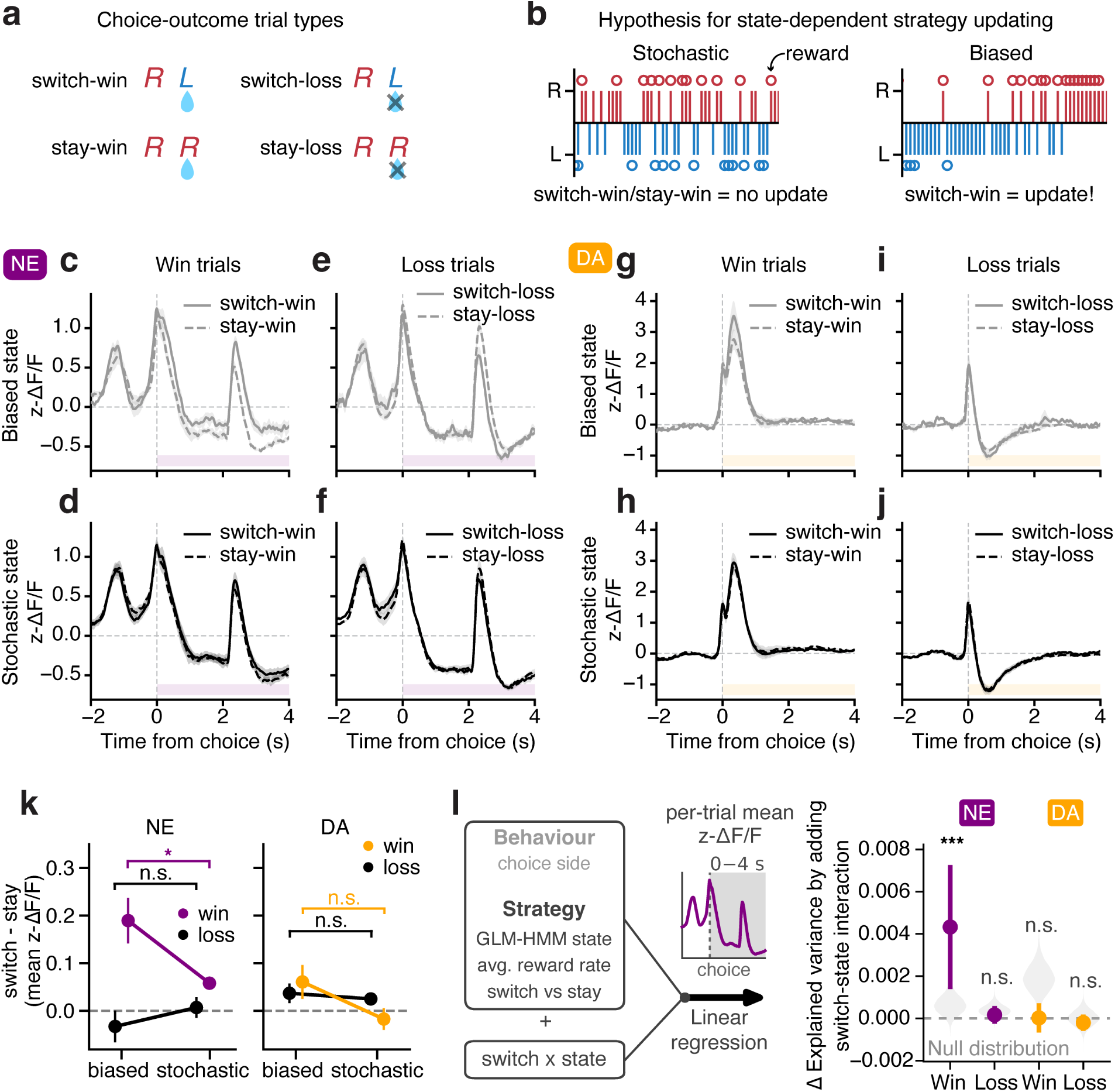
State-dependent updating after rewarded switches. **a)** Schematic showing possible choice (switch/stay) and outcome (win/loss) combinations. **b)** Hypothesis for state-dependent behavioural updating after rewarded switches. **c)** NE activity comparing switch-win (rewarded after switching choice; solid line) and stay-win (rewarded after repeating choice; dashed line) in the biased state. Lines indicate mean *±* SEM across animals (n=5 NE mice). **d)** Same as (c) for the stochastic state. **e)** NE activity comparing switch-loss (unrewarded after switching choice; solid line) and stay-loss (unrewarded after repeating choice; dashed line) in the biased state. **f)** Same as (e) for the stochastic state. **g-j)** Same as (c-f) for DA responses (n=6 mice). **k)** Comparing the average of post-choice (0 – 4 s) NE (left) and DA (right) *switch − stay* activity in the biased versus stochastic state, split by outcome. Data points represent mean *±* SEM error bars across animals. NE switch-stay difference, win trials, stochastic vs. biased: paired t-test across animals, *p* = 0.0358, stochastic mean = 0.0581, biased mean = 0.19; loss trials, *p* = 0.394, stochastic mean = 0.00697, biased mean = -0.0327. DA switch-stay difference, win trials, *p* = 0.0683, stochastic mean = -0.0172, biased mean = 0.0607; loss trials, *p* = 0.739, stochastic mean = 0.0251, biased mean = 0.037. **l)** Difference in explained variance after adding switch*state interaction to the base linear encoding model, fit separately for win and loss trials. Data points represent mean *±* SEM error bars across animals (N = 5 mice for NE, N = 6 mice for DA). Grey region show null distribution obtained by circularly shifting the switch and state labels within each session, and significance was calculated by one-sided permutation test of cohort mean. * p < 0.05, *** p < 0.001, n.s. not significant.

We hypothesised that if NE encodes information relevant for strategy updating, it should therefore distinguish switch from stay trials in the biased state but not in the stochastic state. Indeed, NE activity was elevated following switch-win compared with stay-win trials in the biased state, with the difference emerging after choice and remaining elevated throughout the post-choice period (0–4 s; Figure 4c). This difference was absent in the stochastic state (Figure 4d) and following unrewarded outcomes (Figure 4e,f). Notably, the switch-versus-stay differences were absent for DA responses following either outcome and in either state (Figure 4g-j). We quantified these results for post-choice activity across animals (Figure 4k, Figure S5d–f, m), and by extending our linear encoding model with behaviour and strategy variables to include an interaction between choice switching and behavioural state (Figure 4l). While the comparison between the biased and stochastic states supports a role for NE in representing choice and outcome information relevant for adaptive strategy updating, it should be noted that the biased state does not represent a classic reward-guided strategy. Instead, choices remain shaped by the complex interaction between the animal’s choice history and the opponent’s behaviour. To further understand the difference between NE and DA in shaping strategic choice versus reward-guided learning, we next leveraged a reward-guided task where choice-outcome contingencies provide a more direct and unambiguous basis for updating behaviour.

### Dopamine responses to rewarded switches are amplified in a reward-guided task

We trained the same expert matching pennies mice on a simplified two-armed bandit (2AB) task (Figure 5a; Bari et al., 2019). The high-reward arm always had a non-zero probability of reward, whereas the low-reward arm had zero probability of reward, with these contingencies reversing across blocks without an explicit cue to the animal (Figure 5b). In this task context, reward maximisation depended on identifying and repeatedly selecting the currently more rewarding arm. Animals were first trained on a deterministic version of the task, in which the probability of reward was 1 for the high-reward arm and 0 for the low-reward arm. Under these conditions, reward rate directly reflected the probability of choosing the high-reward arm (Figure 5c). Animals were then progressively trained on more stochastic versions of the task, in which the probability of reward on the high-reward arm was reduced incrementally from 0.9 to 0.6. The resulting reward rates were comparable to those observed during stochastic trials in the matching pennies task (Figure 5d), providing a control for overall reward rate when comparing neuromodulatory responses across tasks. We focused subsequent analyses on the probabilistic 2AB sessions.

**Figure 5.**
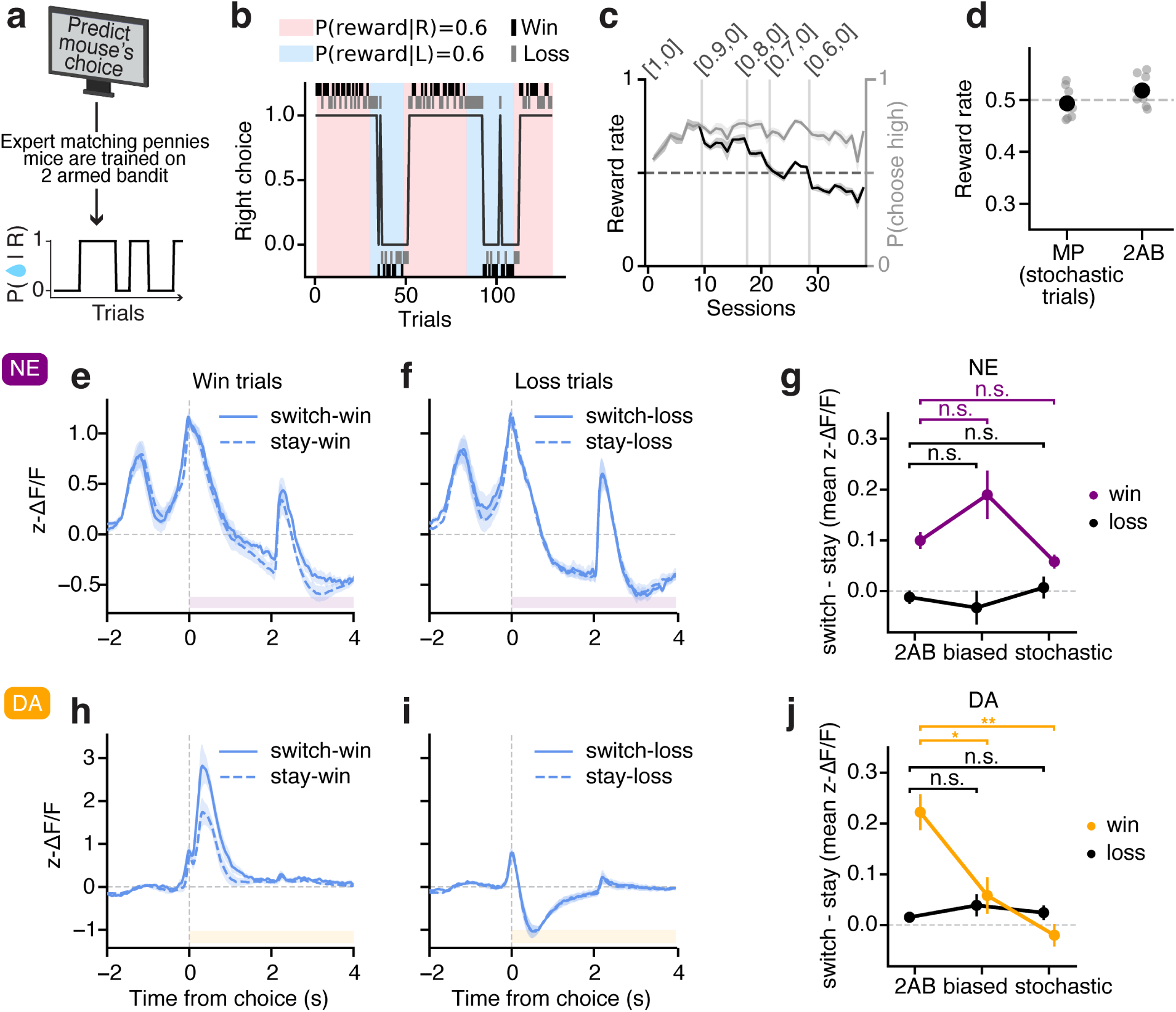
Dopamine exhibits large switch-stay difference only in reward-guided context. **a)** Two-armed bandit (2AB) task schematic. **b)** Example session performance of 2AB task. Red and blue blocks indicate when the high rewarded side is right and left, respectively, black continuous line indicates right choice, and black and grey short lines indicate win and loss trials, respectively. **c)** Reward rate (black) and probability of choosing the high rewarded side (grey) across learning; mean *±* SEM across animals (n = 10 mice). Vertical lines indicate changes in reward contingencies. **d)** Reward rate comparison of all probabilistic 2AB trials versus matching pennies stochastic trials. Black data point indicate mean *±* SEM across mice; grey points show individual mice. **e)** NE activity comparing switch-win (solid line) and stay-win (dashed line) trials in 2AB; mean *±* SEM across animals (n=5 mice). **f)** NE activity comparing switch-loss (solid line) and stay-loss (dashed line) trials in 2AB. **g)** Comparing the average of post-choice (0 – 4 s) NE *switch − stay* activity in 2AB versus biased state trials, and 2AB versus stochastic state trials; mean *±* SEM across mice. NE switch-stay difference, win, 2AB vs. stochastic: paired t-test across animals, *p* = 0.259, 2AB mean = 0.0996, stochastic mean = 0.0581; loss: *p* = 0.595, 2AB mean = -0.012, stochastic mean = 0.00697. NE win, 2AB vs. biased: *p* = 0.241, 2AB mean = 0.0996, biased mean = 0.19; loss: *p* = 0.698, 2AB mean = -0.012, biased mean = -0.0327. **h-j)** same as (e–g) but for DA responses (n=5 mice). DA switch-stay difference, win, 2AB vs. stochastic: paired t-test across animals, *p* = 0.00522, 2AB mean = 0.222, stochastic mean = -0.0387; loss: *p* = 0.115, 2AB mean = 0.0154, stochastic mean = 0.0397. DA switch-stay difference, win, 2AB vs. biased: paired t-test across animals, *p* = 0.0157, 2AB mean = 0.222, biased mean = 0.0351; loss: *p* = 0.615, 2AB mean = 0.0154, biased mean = 0.0251. * p < 0.05, ** p < 0.01, n.s. not significant.

We compared neuromodulatory dynamics across task contexts in a within-subject design, and found that NE showed an intermediate response in the 2AB task relative to those observed in the biased and stochastic states of matching pennies (Figure 5e-g). This intermediate response may reflect differences in task demands (see Discussion). In contrast to NE, DA showed a markedly stronger difference between rewarded switches than stays in the 2AB task compared to the matching pennies task (Figure 5h-j). Unrewarded trials showed little switch-versus-stay difference in either NE or DA (Figure 5f,i). Therefore, when choices were governed by explicit reward contingencies, DA showed a pronounced enhancement following rewarded switches. The strong DA response to rewarded switching in the 2AB task may be consistent with the established role of nucleus accumbens dopamine in encoding reward prediction errors (RPEs) (Day et al., 2007; Hart et al., 2014), such that a rewarded switch generates a positive prediction error when the newly selected arm yields a better-than-expected outcome. To probe that, we next asked how much DA and NE post-choice responses across task contexts could be explained by RPE.

### Reward prediction error accounts for switch-related signals in dopamine, but not in norepinephrine

To obtain a trial-by-trial estimate of reward prediction error (RPE), we fit a Q-learning model following that used in Su et al., 2026, with an additional choice kernel inspired by the work in Wang et al., 2022 (Figure 6a). We found that the model captured mouse behaviour well (Figure 6b) and outperformed previous reinforcement learning models without choice kernels (Lee et al., 2004; Su et al., 2026) in both the 2AB task and the matching pennies task (Figure S8a,b). Interestingly, our 3-state GLM-HMM still outperformed the Q-learning model with a choice kernel in the matching pennies task but not in the 2AB task (Figure S8a, b), suggesting that modelling transitions across discrete behavioural states may be a better description of strategic choices than value-based models.

**Figure 6.**
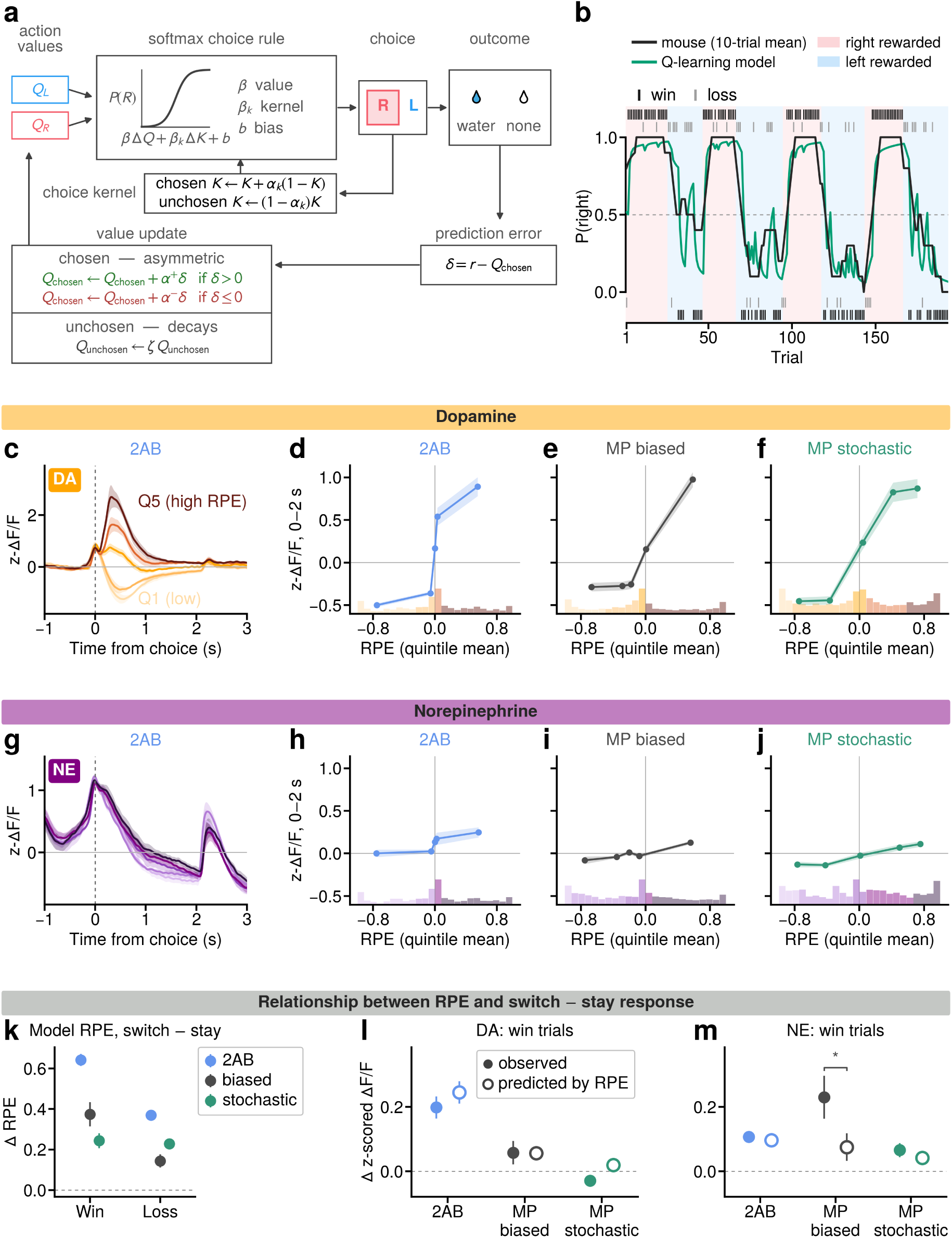
Reward prediction error accounts for switch-related signalling in dopamine, but not in norepinephrine. **a)** Schematic of the Q-learning model fit to mouse behaviour in the two-armed bandit (2AB) and matching pennies task. **b)** Example session fit of choices in the 2AB task. **c)** Choice-aligned dopamine response, during the 2AB task, across mice binned by reward prediction error (RPE) values. Lines show mean and shading shows +/- 1 SEM across mice (n = 5 mice). **d)** Mean dopamine response in the 2AB task (0 – 2 seconds aligned to choice) as a function of reward prediction error. Histogram at the bottom shows the distribution of RPE values pooled across all mice clipped to the 1st – 99th percentile, with bar heights scaled to the square-root of trial counts. **e)** Same as **d** but in the matching pennies task during the biased state. **f)** Same as **d** but in the matching pennies task during the stochastic state. **g)** Same as **c** but for norepinephrine (n = 5 mice). **h)** Same as **d** but for norepinephrine. **i)** Same as **e** but for norepinephrine. **j)** Same as **f** but for norepinephrine. **k)** Difference in RPE between switch trials and stay trials for the model fitted in **a**, in the 2AB task (n = 10 mice), and in the matching pennies task during the biased and stochastic states (n = 11 mice), error bars show +/-1 SEM across models fit to each mouse. **l)** The observed difference in dopamine response between rewarded switch and stay trials for each state and task condition (solid circles), and the predicted difference (open circles) from a linear regression model fit to dopamine response. **m)** Same as **l** but for norepinephrine, observed and predicted difference were compared with a paired t-test across mice. * p < 0.05.

Consistent with previous studies, DA release in the nucleus accumbens tracked RPE in the reward-guided bandit task (Figure 6c, d; Figure S6a, left). Furthermore, we found that the relationship between DA post-win response and RPE generalised to the matching pennies task (Figure 6e, f; Figure S6a, middle and right). We fit a linear mixed-effects model of DA activity as a function of RPE and found no consistent difference in the fitted slopes across animals between task contexts and states (Figure S7a, b). In contrast, while RPE was also encoded by NE (Figure 6g, Figure S6b), it was weaker compared to DA (Figure 6h-j) and relative to other task-relevant representations in NE. Again distinct from DA, the relationship between RPE and NE post-win activity was significantly different across task contexts and states (Figure S7c).

Given the close relationship between DA activity and RPE, we next asked whether the difference in DA activity between rewarded switch and stay trials in the 2AB task (Figure 5e-j) can be accounted for by RPE. We found that model-derived RPE values alone displayed qualitatively similar patterns across switch and stay trials to DA responses, with greater differences during the 2AB task compared to the matching pennies task (Figure 6k). To quantify how much of the switch versus stay activity can be accounted for by RPE, we fit a linear model where the post-choice response on each trial was a function of the RPE and whether the choice was a switch (Methods), and compared the RPE-predicted response with the observed response. For DA, we observed no significant difference between RPE-predicted response and the observed response during rewarded trials (Figure 6l), suggesting that RPE can account for the switch versus stay effect. For NE, RPE-predicted responses accounted for switch versus stay responses during the 2AB task and the stochastic state of the matching pennies task, but not during the biased state of the matching pennies task (Figure 6m). These results further confirm that, unlike DA, the state-dependent NE signals relevant for strategy updating cannot be accounted for by RPE alone.

### Norepinephrine provides trial-by-trial modulation of stochasticity

Within the Q-learning framework, we demonstrated that the primary role of DA release in the nucleus accumbens can be attributed to RPE computation, irrespective of which task or strategy the animal was engaged in. Given the relationship between pre-choice NE activity and strategy-related variables including behavioural state and choice switching (Figure 3d, h), we hypothesised NE may modulate adaptive stochasticity by controlling the temperature and perseverance-related parameters of the softmax choice rule. To test this, we constructed an extension of our best Q-learning model by allowing trial-by-trial NE pre-choice activity to scale the inverse temperature and choice kernel parameters of the model (Figure 7a), and compared its performance in predicting choice to the original model with constant (fit) terms. We found that using NE pre-choice activity to modulate the softmax choice rule significantly improved model performance during the matching pennies task (Figure 7b) but not during the 2AB task (Figure S8c), whereas scaling with DA pre-choice activity did not result in a consistent improvement across animals in either task (Figure 7c, Figure S8c). We also tested whether post-choice NE or DA activity could adaptively modulate learning rate, but did not find a significant model improvement in any condition (Figure S8d). Together, our cross-task comparisons map dissociable functions of NE and DA to distinct computations during adaptive behaviours, with NE primarily controlling choice stochasticity during strategic decision-making and DA primarily guiding reward learning through RPE.

**Figure 7.**
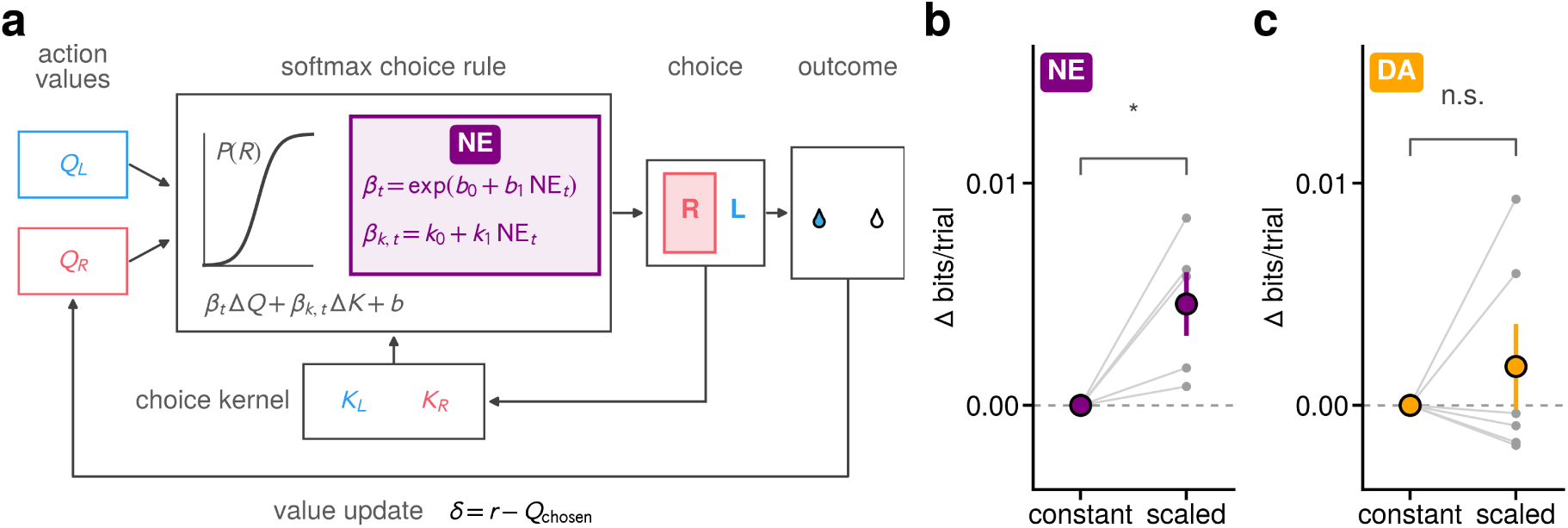
Norepinephrine contributes to modulating temperature and choice kernel in a Q-learning model. **a)** Schematic of a Q-learning model, with the inverse temperature parameter *β_t_* and the choice kernel parameter *β_k,t_* scaled by norepinephrine response on each trial. **b)** Comparison of cross-validated log-likelihood between a model with a constant inverse temperature and choice kernel parameter and a model where norepinephrine scales these parameters per trial. There is a significant difference in cross-validated log-likelihood across mice (p < 0.05, n = 5 mice, one-sample t-test of changes in performance across mice). **c)** Same as **b** but for dopamine (n = 6 mice). * p < 0.05, n.s. not significant.

## Discussion

In this study, we examined how dopamine (DA) and norepinephrine (NE) contributed to strategic choice when the optimal behaviour depended on integrating recent outcomes with longer-term reward and choice information (Figure 1). Combining within-animal comparison of stochastic versus reward-guided choice tasks, recording DA and LC-NE dynamics, and modelling value updating and choice stochasticity, we found distinct neuromodulatory contributions to adaptive behaviours across timescales. DA reliably represented trial-by-trial reward prediction errors (RPEs) across tasks and behavioural states, whereas LC-NE activity contained relatively weaker RPE representations which differed across tasks (Figure 2, 6). NE additionally tracked longer-timescale reward and strategy-related information (Figure 3), even when controlling for movement, learning, and task engagement (Figure S4, Figure S5); and its relationship with choice and outcome depended on the animal’s behavioural strategy (Figure 4, 5). Lastly, using trial-by-trial NE activity to modulate the noise of choice generation improved performance of a reinforcement learning model (Figure 6, 7), providing a computational account linking NE activity to adaptive stochasticity. These findings suggest that although both neuromodulatory systems carry teaching signals that guide learning and choice, NE additionally reflects arousal and broader behavioural context to shape strategies in complex decision environments.

The most robust feature of nucleus accumbens DA release across our experiments was its relationship with RPE, consistent with the extensive literature linking phasic DA signalling to temporal-difference RPEs (Bayer and Glimcher, 2005; Cohen et al., 2012; D’Ardenne et al., 2008; Montague, Dayan, and Sejnowski, 1996; Schultz, Dayan, and Montague, 1997). Furthermore, we found that DA’s relationship with RPE was reliable across tasks despite substantial differences in reward statistics and strategic demands (Figure 6). It should be noted that our experiments were not designed to test other proposed functions of DA such as distributional value representation (Dabney et al., 2020), motivational drive (Howe et al., 2013), vigour (Hamilos et al., 2025; Niv et al., 2007), action selection (Gershman et al., 2024), value-free learning (Greenstreet et al., 2025), retrospective learning of cue-outcome associations (Burke et al., 2026; Jeong et al., 2022). Nevertheless, our findings validate that RPE provides a strong and consistent description of DA during the outcome period, and it is important to consider whether other observed representations, such as switch-versus-stay selectivity (Figure 5), transient state encoding (Figure 3g) and reward rate signals (Figure S3e, right), can all be accounted for by RPE.

While LC-NE activity also encoded RPE, the relationship between NE and RPE was less consistent than the corresponding DA-RPE relationship (Figure 6). In a recent study, anatomical and physiological characterisation of LC-NE neurons revealed functional heterogeneity within the nucleus, identifying a population of cortex-projecting dorsal LC neurons whose activity increases with RPE (Type I), and ventral LC populations that responded to the absence of reward (Su et al., 2026). Although our fibre photometry recordings did not differentiate these subpopulations, the observed NE responses were consistent with the Type I LC-NE neurons that scaled with RPE in all mice (Figure S6b). It remains to be investigated whether the seemingly weaker RPE representation in our LC-NE recordings can be attributed, at least in part, to averaging across heterogeneous neuron types.

In contrast, LC-NE activity reliably encoded salient task events as well as strategy-related variables (Figure 2, Figure 3), both of which may be related to LC as a global model failure system (Jordan, 2024). We found a strong correlation between LC-NE activity, pupil diameter, and salient events (Figure 1i, Figure S2), supporting a large literature linking both NE and pupil fluctuations to changes in arousal and internal state (Costa and Rudebeck, 2016; Joshi et al., 2016; Reimer et al., 2016), in coordination with other neuromodulatory systems (Cazettes et al., 2021; Khalighinejad and Rushworth, 2026). However, by simultaneously recording LC-NE activity and pupil responses, we revealed that NE carried unique task-relevant information that was not explained by pupil (Figure S2k,l). Therefore, our results highlight the value of directly measuring neuromodulatory activity when associating cognitive functions to arousal-related signals, and caution against using pupil diameter as a direct readout of LC-NE activity (Megemont, McBurney-Lin, and Yang, 2022; Reimer et al., 2016).

Between phasic representation of salient events, NE activity also encoded longer-term strategy-related information, before choice and during outcome evaluation. Pre-choice NE signals varied with average reward rate, switching of upcoming choice and behavioural state inferred from the GLM-HMM (Figure 3). While average reward rate and probability of choice switching were both correlated with the behavioural state, we confirmed that NEs relationship with these variables persisted within each state (Figure S3f,g). These results are consistent with previous studies where optogenetic and chemogenetic elevation of frontal NE release promoted stochastic behaviour (Tervo et al., 2014; Wang et al., 2025). Furthermore, in a reinforcement learning framework, scaling the parameters governing choice stochasticity with trial-by-trial pre-choice NE activity improved model performance for the stochastic but not the reward-guided task (Figure 7). This finding supports a meta-learning framework where NE regulates the inverse temperature, allowing the balance between exploitation and exploration to adapt to changing environmental demands (Doya, 2002), which may be especially important for competitive contexts.

On the other hand, post-choice NE activity was selectively enhanced for rewarded choice switches in behavioural states containing structured choice patterns (Figure 4), and these signals cannot be accounted for by RPE (Figure 6). We hypothesise that the state-dependent NE signal for switching may be related to serial hypothesis testing before animals reached the optimal strategy of choosing left and right randomly. In other words, while pre-choice NE activity regulates strategic choice stochasticity, NE outcome activity contains information that guides the decision to maintain or revise its current strategy. Because hypothesis testing may be reduced in simple reward-guided contexts and suppressed during “random" choice generation, these neural signatures were weakened in the two-armed bandit task and the stochastic state of the matching pennies task. This interpretation more generally supports theories in which LC-NE contributes to flexibility and the regulation of learning according to environmental stability (Aston-Jones and Cohen, 2005a; Doya, 2002; Yu and Dayan, 2005). In conventional reward-guided tasks, an unexpected reward or omission both generate an RPE and indicate that animals should change their behaviour. Competitive environments weaken this coupling, allowing the relationship between neuromodulatory activity and strategy updating to be examined separately from teaching signals. By comparing DA and NE dynamics in both competitive and reward-guided task contexts, we revealed complementary neuromodulatory mechanisms for fast timescale reward learning and slower timescale strategy formation that maximises long-term benefits in interactive decision environments.

## Methods

### Animals

All experiments were performed in accordance with the UK Home Office regulations (Animals Scientific Procedure Act 1986, PPL: PP4459082) following ethical approval by Sainsbury Wellcome Centre Animal Welfare Ethical Review Body. A total of 11 mice (5 female, 6 male) 8–16 weeks old at the start of experiments were used in this study. All 11 mice provided fibre photometry and behavioural data. 5 Dbh-Cre mice were used for norepinephrine recordings and 6 C57BL/6J wild type (from Charles River Laboratories) mice were used for dopamine recordings. Experiments were carried out in the dark phase of the 12/12 h light/dark cycle.

### Surgery

Surgical procedures were carried out within aseptic guidelines. All animals underwent headplate implantation surgery. Mice were induced under anaesthesia using isoflurane (4% induction, 1.5% maintenance) and were given a subcutaneous injection of analgesic (Metacam). The scalp was shaved and cleaned using an antimicrobial skin cleanser (Hibiscrub). Mice were secured onto a stereotaxic frame and placed over a heat pad at 37*^◦^*C. Temperature was monitored using a rectal probe and eye lubricant was used to protect their eyes. The scalp was cut and removed to expose the surface of the skull, which was cleaned and dried using saline. After induction under 4% isoflurane, animals underwent a craniotomy, stereotactic virus injection and fibre placement at the injection site. Dbh-Cre mice used to record LC-NE activity received an injection of GCaMP8s (AAV-hSyn-flex-GCaMP8s) at coordinates targeting the LC (in mm: AP -5.4, ML *±*0.85, DV -3.68). Optic fibre cannulas (Doric, 400 ţm 0.37 NA, MFC_400/470-0.37_4.0mm_ZF1.25(G)_TP or MFC_400/470-0.37_4.0mm_ZF1.25(G)_FLT) were implanted at the same coordinates as the injection site. Wild type mice used for DA recordings in NAc were injected with dLight1.1 (AAV5-CAG-dLight1.1-WPRE-SV40) at coordinates AP +1.05, ML *±*1.6, DV -3.85 from brain surface (in mm). Optic fibre cannulas (Doric, 400ţm 0.37NA, MFC_400/470-0.37_5.0mm_ZF1.25(G)_TP) were implanted at the same coordinates as the injection site. A headplate was attached using MetaBond and dental cement.

### Behaviour

#### Hardware

Behavioural training was conducted in custom-built aluminium boxes (50*×*50*×*50 cm) lined with sound-attenuating foam, using the apparatus and behavioural acquisition procedures described previously (Aloor et al., 2026; Gauld et al., 2026). Briefly, mice were head-fixed by clamping the headplate bilaterally between two metal arms, with the body supported in an acrylic tube that allowed free adjustment of posture. Auditory stimuli were delivered through a front-facing electrostatic speaker (ED1 with ES1 driver; Tucker-Davis Technologies). Two lick ports (10 cm lengths of 1/16*^′′^ ×* 1/32*^′′^* steel tubing; Coopers Needle Works) were gravity-fed with water through solenoid pinch valves (225P011-11; NResearch) mounted outside the chamber. The unitary reward volume (1 *µ*l) was calibrated by weight and matched across ports and rigs. Rewards larger than 1 *µ*l were delivered as multiple pulses. Licks were detected with a dual-channel comparator circuit adapted from a Janelia Research Campus design (https://github.com/janelia-experimental-technology/Comparator-Dual-Lick-Detector). The lick ports were held in a 3D-printed mount driven by two servo motors (XL-320; Robotis), allowing motorised translation in the anterior–posterior and mediallateral axes, with manual adjustment in the vertical axis. Anterior–posterior movement of the ports marked the opening and closing of the response window. Two infrared-sensitive cameras (Raspberry Pi High Quality Camera with 8–50 mm C-mount zoom lens) provided frontal and lateral views for recording pupil and orofacial movements, with the chamber illuminated by an infrared LED array and ambient white light. Video was acquired at 90 fps by a dedicated Raspberry Pi 4B per camera (https://github.com/larsrollik/rpi_camera_colony), which also logged timestamped TTL pulses to align frames to trial onset.

#### Matching Pennies computer opponent

The Matching Pennies computer opponent and behavioural task were implemented as described previously (Aloor et al., 2026; Barraclough, Conroy, and Lee, 2004; Tervo et al., 2014). Briefly, the opponent predicted the animal’s upcoming choice from statistical regularities in its previous choice and reward history. The algorithm tracked choice-only and choice-reward sequences of lengths 1–4, together with overall choice bias, and evaluated each feature using a binomial test against the null hypothesis of equal left and right choice probability (*P* = 0.5). When at least one feature reached the significance threshold (*p <* 0.05), the prediction was based on the feature providing the strongest statistical evidence; otherwise, the opponent selected left or right with equal probability.

#### Matching pennies task

Mice were habituated to head fixation for 3–5 days before undergoing two shaping phases to establish reliable licking and alternating left-right choices (Aloor et al., 2026). In the first phase, mice learned to associate licking with reward, receiving 3 ţl of water from the spout they licked. To prevent the development of a side preference, a closed-loop system monitored licking and, when *≥*80% of licks over 20 trials were directed to one side, the lick port was automatically displaced laterally by 0.5 mm in the direction that counteracted the bias. In the second stage, mice learned to sample both spouts. Only one side was rewarded at a time, and the rewarded side switched once the animal had chosen it on three trials. In this stage the spouts were retracted at trial onset and advanced into licking range only during the choice period. Animals were subsequently trained on the Matching Pennies task, in which each trial began with a 10 kHz auditory cue followed by a variable delay of 1–1.5 s, drawn from a uniform distribution, before the lick ports were extended. Mice needed to make a choice within a 2 s response window, after which reward was either delivered immediately if the choice was not predicted by the computer opponent or not delivered at all when the computer prediction was correct. The ports remained accessible for 2 s, allowing for reward consumption before being retracted. The inter-trial-interval was drawn from a truncated exponential distribution of 5–8 s.

#### Two-armed bandit

Following Matching Pennies training, animals were trained on a two-armed bandit task in which the left and right lick ports were associated with different reward probabilities that changed across uncued blocks. Block size was determined by a Poisson distribution with *λ* = 30. The trial structure was designed to match the same sensorimotor cues as Matching Pennies. Trial start, indicated by a 10 kHz auditory cue, was followed by a variable delay of 1–1.5 s. Lick ports were extended to initiate the start of the response window. Animals had 2 s to make their choice by licking left or right, with reward delivered immediately to the rewarded port on correct trials. Trials were separated by a randomly selected inter-trial-interval of 5–8 s. The first stage of training used a deterministic reversal contingency, in which one port had a reward probability of 1 and the other had a reward probability of 0. Once animals achieved approximately 70% choice accuracy for the higher-probability port across 2–3 sessions, the high-rewarded probability was progressively reduced across training stages from 0.9 to 0.6, while the alternative port remained unrewarded (*P* = 0).

### Behaviour analysis

#### Exponential smoothing kernel

Within-session reward rate and choice probabilities were estimated using an exponential smoothing kernel:

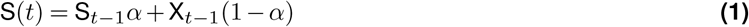

where *X_t_* is the trial-by-trial measure (e.g., reward or choice on trial *t*) and *α* = 0.95. This is equivalent to convolving the trial history with an exponentially decaying kernel, in which the weight given to a trial *k* trials in the past is (1 *− α*)*α^k^*, giving a time constant of *τ* = *−*1*/* ln *α ≈* 20 trials. Smoothing was applied recursively across the full session, with *S*_0_ initialised to the mean of the session.

#### Shannon’s entropy

Choice sequence variability was quantified using Shannon entropy calculated over five-choice sequences comprising the current choice, the two preceding animal choices, and the opponent’s two preceding choices.

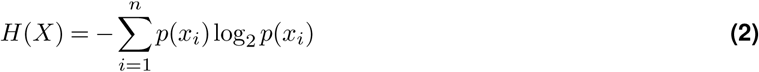

### Reaction time analysis

Reaction time is defined as the time difference between port extension and the time of choice defined from the animal’s first lick. Missed trials were excluded from the reaction time analysis. For Figure S2b-f, reaction time was re-scaled to the percentage change from each session’s own median:

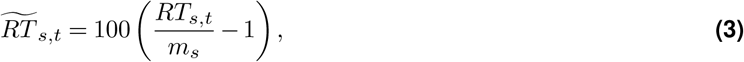

where RT*_s,t_* is the reaction time on trial *t* during session *s*, *m_s_* is the median reaction time in session *s*. To model the variables contributing to reaction time in Figure S2g-h, we log-transformed the reaction time within session and obtained the z-scored values. To dissociate the contributions of behavioural variables on the reaction time of each trial, we fit a linear regression on the log-transformed reaction time either only as a function of the post-outcome response in the previous trial, or additionally as a function of reward in the previous trial, reward in the current trial, trial index, and log-transformed reaction time in the previous trial (Figure S2g). We then compared the obtained coefficients on those two regression models (Figure S2h, and performed one-sample two-sided t-test across mice to test for significant difference from zero (n = 5 mice for norepinephrine, n = 6 mice for dopamine, n = 9 mice for pupil).

### GLM-HMM

A GLM-HMM (Ashwood et al., 2022) was fit to mouse choice data to identify latent behavioural states. The model included three regressors describing general side bias, the previous rewarded choice, and the previous unrewarded choice. Parameters were estimated by maximum likelihood using the expectation-maximisation algorithm. Models containing 1–5 states were compared using leave-one-session-out cross-validation, with ten random initialisations per fold and test log-likelihood normalised to the corresponding 1-state model (Aloor et al., 2026). State labels and GLM parameters were aligned across model initialisations and cross-validation folds using constrained k-means clustering, and median state posterior probabilities across initialisations were used for subsequent analyses. Trials were assigned to a state when its posterior probability exceeded 0.8.

### Pupil preprocessing

Pupil size was measured from side-camera videos recorded at 90Hz during each behavioural session. Pupil keypoints were tracked using SLEAP (Pereira et al., 2022). 8 keypoints were defined around the pupil margin, and approximately 500 frames were labelled by hand in the SLEAP graphical interface. These frames were used to train a single-instance model. The trained model was then used to predict keypoint locations on every frame of all remaining videos. For each session, the tracked keypoint coordinates were converted into a frame-by-frame estimate of pupil area. Keypoints that that appeared as outliers from the rest of the pupil outline were removed first. For each frame, we computed the centroid of all keypoints and each keypoint’s Euclidean distance from it. Any keypoint whose distance exceeded the mean distance in that frame by more than 2.5 standard deviations was removed An ellipse was then fitted to the keypoints by least squares (Hammel and Sullivan-Molina, 2020), and pupil area was computed as *A* = *πab*, where *a* and *b* are the semi-major and semi-minor axes of the fitted ellipse. Frames with missing keypoints or failed ellipse fitting were not included in analysis. The pupil-area time series was low-pass filtered (4 Hz cutoff). Extreme values were then removed with a median absolute deviation (MAD) criterion. Frames with filtered area below *Ã−* 3 MAD or above *Ã*+ 10 MAD, where *Ã* is the session median, were excluded. Finally, the trace was z-scored within session using the mean and standard deviation of the remaining valid frames. Excluded frames were not interpolated. Camera frames were aligned to behaviour using TTL pulses on each trial. Each pulse was assigned to the first camera frame whose timestamp was at or after the pulse. The z-scored pupil trace was then epoched from *−*3 to +5 s relative to the first lick.

### Face movement analysis

To quantify the relationship between uninstructed facial movement and neuromodulatory signals (Figure S4), we filmed the face of the mouse from the side, we captured movement from the eye, whiskers, snout, and in some cases paw movement (Figure S4a). The video data was downsampled to 160 *×* 120 and grayscaled, and motion energy was defined as the mean absolute gray-level difference of consecutive frames (Figure S4b-c). During this calculation, we masked the spout and mouth area of the video, so that port extension and retraction movement were excluded from the quantification of mouse movement. Motion energy traces were z-scored within each session (Figure S4d). To correlate face motion and pre-choice neuromodulatory signal (Figure S4e), we computed the mean motion energy from -2 to 0 seconds before the choice time on each trial. For each mouse, we then divided this mean motion energy value into ten bins, and correlated the mean value within each bin and the pre-choice norepinephrine or dopamine signal corresponding to the same trials. To compute the cross-correlation between neuromodulatory signals and facial motion energy (Figure S4f), we downsampled both signals to 30 Hz. We define “event-free" periods as time periods that is at least 3 seconds from the sound cue or time of choice. To investigate whether there were state differences in pre-choice neuromodulatory signals after accounting for facial movement (Figure S4g), for each mouse, we fit an ordinary least-square regression model where the pre-choice neuromodulatory signal was the target and the regressors include an intercept per session, an indicator variable denoting the state (stochastic or biased), and the pre-choice motion energy. After fitting this model, we subtracted the motion contribution to the neuromodulatory signal, then regressed the neuromodulatory signal as a function of the state variable across sessions to obtain a slope value that excluded the effect of movement. Finally, we performed a one-sample t-test across mice to test for a significant difference from 0. We repeated the same analysis using the reward rate as a regressor instead of state (Figure S4h).

### Fibre photometry

#### Hardware and data acquisition

Fibre photometry recordings were performed using a dual-wavelength excitation system. Excitation light at 465 nm, corresponding to the calcium-or dopamine-sensitive channel, and 405 nm, corresponding to the isosbestic control channel, was delivered using connectorised LEDs (Doric Lenses). Fluorescence was collected through a four-port fluorescence mini cube with an integrated photodetector (Doric Lenses) and coupled to the implanted fibre optic cannula using low-autofluorescence optical fibre patch cords (400 *µ*m diameter, 0.37 NA; Doric Lenses). Fluorescence was acquired at 120 fps using a pyPhotometry acquisition board (Akam and Walton, 2019), which additionally controlled LED excitation. Behavioural event timing was synchronised with photometry recordings by sending TTL pulses from the Bpod behavioural control system to the digital input of the pyPhotometry board. The pyPhotometry software alternated between 465 and 405 nm excitation on successive frames throughout each recording. LED power was set to the minimum level required to reliably detect fluorescence, typically around 5 *µ*W at the fibre tip, and was adjusted across training as necessary to maintain signal quality while minimising photobleaching and phototoxicity.

#### Data preprocessing

Raw 465 nm and 405 nm fluorescence signals were first denoised using a median filter followed by a 10 Hz low-pass filter. Within-session photobleaching was corrected by fitting and subtracting a linear trend from each signal, followed by removal of residual slow baseline drift using the moving 5th percentile calculated within a 60 s sliding window. The 405 nm isosbestic signal was then linearly regressed against the corresponding 465 nm signal to account for signal fluctuations unrelated to the calcium-or dopamine-dependent fluorescence, including movement- and hemodynamic-related artefacts. The corrected 465 nm signal was expressed as Δ*F/F*_0_, with *F*_0_ defined by the fitted baseline used for bleaching correction. Finally, Δ*F/F*_0_ traces were z-scored within each session to facilitate comparison across animals and sessions.

### Linear encoding model

To dissociate the contributions of task and behavioural variables on neuromodulatory activity, we fit regression models to the activity. For each mouse, we took the fluorescence (ΔF/F) activity aligned from -1 second to 7 seconds relative to sound onset, resampled at 30 Hz, and concatenated trials to one pseudo-continuous time series, which was then z-scored. This gives us the target variable *Y*. The design matrix *X* was constructed via a set of Toeplitz matrices with one column per lag bin around the event of interest. These events included the sound onset, port extension and port retraction, all of which had a support of 0 to 1 second aligned to their onset, and first lick onset, lick direction, reward, GLM-HMM state, switch/stay, and smoothed reward rate, all of which had a support of -2 to 4 seconds aligned to lick onset. Reward and lick direction were +1/-1 coded, and other variables were 0/1 coded. For GLM-HMM state coding, the left-biased and right-biased state was combined to form a single biased state. The feature matrix was z-scored per column during the fitting process. We used a ridge regression model with a fixed hyperparameter *λ* = 1.0, and the model was evaluated using 5-fold cross-validation across trials, so that all time points of a trial fall in the same fold. We calculated the cross-validated explained variance to compare across models.

To calculate the critical time points where the strategy term (GLM-HMM state, average reward rate, and choice switch) contributed additional explained variance (Figure 3i), we re-aligned held-out predictions from sound onset to time of first lick from -2 seconds to 4 seconds after the first lick. Explained variance was then recomputed for each choice-aligned time bin across trials.

To test for a reward- and state-dependent effect of switch versus stay response in norepinephrine and dopamine (Figure 4k, l), we first computed the mean z-scored activity of dopamine and norepinephrine ΔF/F signals during the post-choice period (0 - 4 seconds aligned to choice) for each trial. For win and loss trials, we fit two separate models each to model the neuromodulatory activity *y*. The interaction model tested is given by:

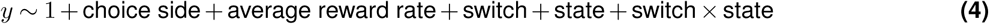

and was tested against a baseline model where the final interaction term between switch and state was removed. The model was fit using ordinary least squares and we compared the 5-fold cross-validated explained variance between the interaction model and the baseline model. The null distribution for each win/loss condition and neuromodulator was created by first circularly shifting the switch and state condition of trials per session and generating the interaction value from the shifted labels, then fitting a model using this shifted interaction label with the other regressors kept intact. This process was repeated 500 times per mouse, and the cohort-level null distribution was created by performing 20,000 repeated draws of one random sample per mouse, then averaging across mice.

The linear encoding analyses related to pupil (Figure S2i–l) followed the same methods as that described in the earlier regression methods, but using z-scored pupil activity as the target. The photometry signal was resampled at 30 Hz to match the sampling rate of the pupil signal. The pupil dataset consisted of 5 mice from the NE cohort (JOA-M-0022–0026) and 4 from the DA cohort (JOA-M-0020, -0027, -0028, -0036). For regression of dopamine and norepinephrine activity using pupil as a regressor, we included 20 time-shifted copies of the pupil trace over a 5 second range in steps of 0.263 seconds, shifting from time *t* to time *t* + 5 seconds, based on the observation (Figure 1i) that norepinephrine activity preceded the pupil response. For linear encoding models involving only pupil as the target (Figure S2i, j), 154 sessions from these 9 mice were used. For linear encoding models involving dopamine and norepinephrine as the target and pupil as a regressor, 91 sessions from these 9 mice were used.

### Linear regression predicting mean activity from reward rate and state

To test whether trial-by-trial fluctuations in reward rate were associated with neuromodulator activity, and whether this relationship depended on behavioural state, we fit separate linear regressions for three different trial epochs—pre-choice (*−*2 to 0 s relative to choice), post-choice (0 to 2 s), and post-outcome (2 to 4 s)—and for each neuromodulator group (NE, *n* = 5; DA, *n* = 6). For each trial, activity was defined as the mean *z*-scored Δ*F/F* signal within the epoch window. Reward rate was the smoothed trial-wise reward rate, and behavioral state (stochastic vs. biased) was assigned from the fitted GLM-HMM. Trials without a clear state assignment, *p*(state) *≤* 0.8, were excluded from all state-dependent models.

For each subject and epoch, we fit the following ordinary least-squares model:

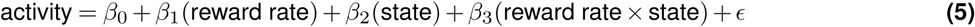

State was effect-coded (*−*1 = biased, +1 = stochastic), and reward rate was mean-centered within each subject’s trials for that epoch prior to forming the interaction term, so that *β*_1_ reflects the reward-rate slope at each subject’s own mean reward rate rather than at reward rate = 0. Models were fit independently per subject given the small number of subjects per group.

For each term (*β*_1_, *β*_2_, *β*_3_), the resulting per-subject coefficients were tested against zero across subjects with a one-sample *t*-test. *p*-values were corrected for multiple comparisons across all three terms and three epochs within each group (9 tests) using the Benjamini-Hochberg false discovery rate (FDR) procedure (*q <* 0.05 considered significant).

### Logistic regression predicting switch from pre-choice activity

To test whether neuromodulator activity preceding a choice predicted whether the animal would switch its choice on the upcoming trial, we fit a per-subject logistic regression:

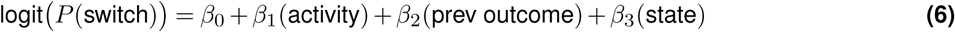

where “switch” indicated whether trial *t*’s choice differed from trial *t −* 1’s choice, activity was the mean *z*-scored Δ*F/F* signal in the pre-choice epoch (*−*2 to 0 s relative to choice), and prev_outcome was the reward outcome (win/loss) of trial *t−* 1, included as a control covariate. Trials were excluded if either the current or preceding trial was a miss trial, or if the trial was the first of its session.

Models were fit separately per subject via logistic regression. The resulting per-subject activity coefficients (*β*_1_ in Equation 6) were tested against zero across subjects within each group (NE, *n* = 5; DA, *n* = 6) using a one-sample *t*-test.

### Within-session and session-position controls

Stochastic state occupancy generally increases as animals learn the matching pennies task, meaning differences in neuromodulatory signal between states may be attributed to differences across days that is not due to state differences. To test this, we compared the results obtained via pooling stochastic and biased state trials from all trials across sessions and computing their difference, with a separate approach computing the difference within each session followed by averaging across sessions (Figure S5a). Sessions were only included in the within-session analyses if they had at least 20 biased and 20 stochastic trials. Using both methods, we compared the difference in norepinephrine signal during the biased and stochastic state (Figure S5b), and fit two ordinary least square (OLS) regression models that model the pre-choice (-2 to 0 seconds aligned to choice) norepinephrine activity as a function of state (stochastic or biased) and reward rate, without (pooled model) and with (within model) an intercept per session. We then performed a one-sample t-test across mice to test for a significant difference of the state coefficient from 0 (Figure S5c). Using the same approach, we compared the difference in norepinephrine signal (0 - 4 seconds after choice) in switch versus stay trials either over all sessions (pooled method) or first computing this difference within each session, followed by averaging across sessions (within method). For this analysis, only sessions with at least 3 trials in every combination of state and switch versus stay condition were included. Statistical significance was tested by performing paired t-tests in stochastic versus biased trials across mice for win and loss condition separately.

State occupancy and behavioural proxies of engagement including reaction time, miss rate and number of licks can vary as a function of trials within a session (Figure S5g-j). To control for this whilst comparing differences in neuromodulatory signals across states, for each mouse, we first divided trials into 10 bins within a session, and within each bin, we subsampled trials from each state to match the trial number of the state with fewer trials (Figure S5k). We repeated this subsampling process 20 times and reported the mean values. The unmatched results used random subsamples of the same size so that the comparison between matched and unmatched results cannot be attributed to differences in trial counts. Using this procedure, we repeated the pre-choice OLS regression analysis to test for a difference in norepinephrine signal between states (Figure S5l), and for differences in post-choice signal during switch versus stay trials (Figure S5m).

### Q-learning model

To quantify the relationship between the neuromodulatory signals and reward prediction error (RPE), we first fit a Q-learning model to the behaviour. The model was based on Su et al., 2026, with a choice kernel following that of Wang et al., 2022. On each trial, the model makes a choice based on the equation

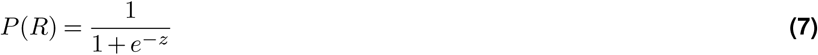

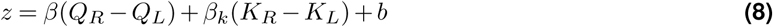

where *Q_L_* and *Q_R_* were the values for left and right choices, *β* was the inverse temperature parameter, *b* was the bias term, *K_L_* and *K_R_* were the choice kernels for left and right choices, and *β_k_* was the choice kernel weight.

After each choice *c_t_ ∈ {L, R}* on trial *t* and reward feedback *r_t_ ∈ {*0, 1*}*, the chosen value was updated via

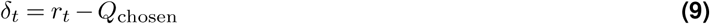

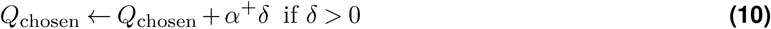

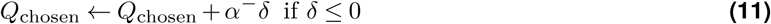

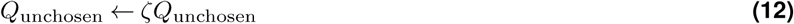

where *δ_t_* was the reward prediction error, *r_t_* was the reward on trial *t*, *Q*_chosen_ was the chosen Q-value on trial *t*, *α*^+^ was the learning rate for rewarded choices, *α^−^* was the learning rate for unrewarded choices, *Q*_unchosen_ was the Q-value of the choice not chosen on trial *t*, and *ζ* was a decay term for the unchosen value. The choice kernel was reward-independent and updated via

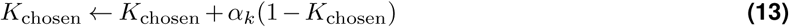

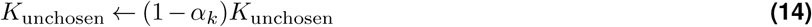

where *K*_chosen_ was the choice kernel for the choice chosen on trial *t*, and *α_k_* sets the update rate of the choice kernel.

For each mouse, we fit a Q-learning model across all trials and sessions separately for the two-armed bandit and matching pennies task. The model contained 7 free parameters: *α*^+^*, α^−^, ζ, β, b, α_k_, β_k_*, with bounds *α*^+^ *∈* (0, 1)*, α^−^ ∈* (0, 1)*, α_k_ ∈* (0, 1)*, ζ ∈* (0, 1)*, β ∈* [10*^−^*^3^, 20]*, β_k_ ∈* [*−*20, 20]*, b ∈* [*−*5, 5]. The Q-values *Q_L_, Q_R_* and choice kernels *K_L_, K_R_* were initialised to 0 and reset at the beginning of each session. Miss trials were excluded from analysis. The model was fit via maximum likelihood using the L-BFGS-B algorithm. This procedure was repeated four times, one from fixed initial values, and four from a random draw within bounds, and the best solution was kept and evaluated on the test set.

For model comparison, we performed 5-fold cross-validation, where each fold contained an equal number of trials from each session. The same cross-validation folds were used to compare the performance between the Q-learning model with choice kernel, GLM-HMM with 3 states, Q-learning model from Lee et al., 2004, and the Q-learning model from Su et al., 2026 (Figure S8a, b). Latent variables (Q-values, choice kernels, GLM-HMM state) were propagated across all trials and the likelihood evaluated on held-out trials. We use held-out Bernoulli log-likelihood in bits per trial as the evaluation metric.

To extend the model to allow norepinephrine to scale the temperature and choice kernel (Figure S8c), we fit the following two link functions:

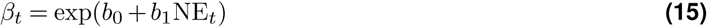

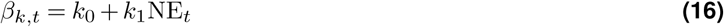

where NE*_t_* esd the mean within-session z-scored Δ*F/F* of norepinephrine activity -1 to 0 seconds relative to the time of choice, and *b*_0_*, b*_1_*, k*_0_*, k*_1_ were free parameters. The inverse temperature *β_t_* was first linearly scaled by NE*_t_* followed by an exponentiation such that the values stay positive, and the choice kernel scaler *β_k,t_* was linearly scaled such that positive values promote perseveration and negative values promote alternation. We repeated the same analysis with dopamine by fitting separate link functions.

To extend the model to allow norepinephrine and dopamine to scale the learning rate (Figure S8d), we fit the following two link functions:

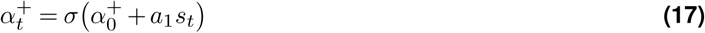

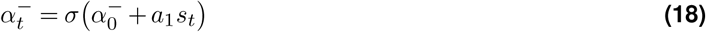

where *σ* was the sigmoid function, *s_t_* was the mean post-outcome activity (2 - 4 seconds after choice) of either norepinephrine or dopamine, and *α*_0_^+^, *α*_0_^-^, *a*_1_ are free parameters.

To compare the relationship of dopamine and norepinephrine with RPE across reward outcome, task and states (Figure S7a-c), we first obtained the intersection of the 5th to 95th percentile of RPE values across the two-armed bandit task, the matching pennies task in the biased state, and the matching pennies task in the stochastic state. This ensured that the same RPE range was used across conditions. For each mouse, we fit an ordinary least squares regression of the post-choice signal (0 - 2 seconds after choice) against the RPE, and obtained the slope. To test for a difference in slope across task and state conditions, we performed a likelihood-ratio test between a linear mixed-effects model where the slope depends on the condition and a random effect of the mouse identity, and a model with only a random effect of the mouse identity. For pairwise comparison across each pair of conditions, we performed paired t-test across mice.

To test whether RPE could account for the difference in post-choice (0.2 – 4 seconds after choice) activity between switch versus stay trials across tasks and states (Figure 6k–m), for each mouse, task, state and reward condition, we fit a linear model via ordinary least squares:

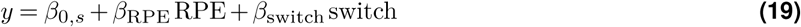

where *y* was the observed post-choice response, *β*_0_*_,s_* was an intercept term for each session *s*, *β*_RPE_ captured the relationship between the neuromodulatory signal and RPE, and *β*_switch_ captured the difference in activity between switch and stay trials. We then obtained a residual term by comparing the observed neuromodulatory signal with the prediction made using the model based on the RPE and intercept terms only, and tested whether this residual was significantly different from zero using a one-sample t-test (Figure 6l, m).

## DATA AND CODE AVAILABILITY

Code and data related to this manuscript are available at:

https://gin.g-node.org/DELab/matching-pennies-neuromodulation-2026

## ACKNOWLEDGEMENTS

This work was funded by the Sainsbury Wellcome Centre Core Grant from the Gatsby Charitable Foundation (GAT4057) and Wellcome (318818/Z/24/Z), by a UK Research and Innovation grant (EP/Y008804/1); and by the Gatsby Initiative in Brain Development and Psychiatry (GAT3955). We thank members of the Duan laboratory and members of the Erlich laboratory for discussion and advice; D. Lee, O. Gauld, C. Sun, and M. Stephenson-Jones for comments on the manuscript; SWC Neurobiological Research Facility for animal support; and FabLabs for contributions to hardware design.

## AUTHOR CONTRIBUTIONS

C.A.D. and J.A. conceived the project. J.A. and T.P.H.S. performed surgeries. J.A., J.G. and J.O.T. contributed to mouse behavioural training and fibre photometry experiments. J.A. and T.P.H.S. analysed data, with contributions from J.G. on pilot analyses. C.A.D. provided supervision and input to all data analysis. C.A.D., J.A. and T.P.H.S. wrote the manuscript.

## Supplementary Information

**Figure S1.**
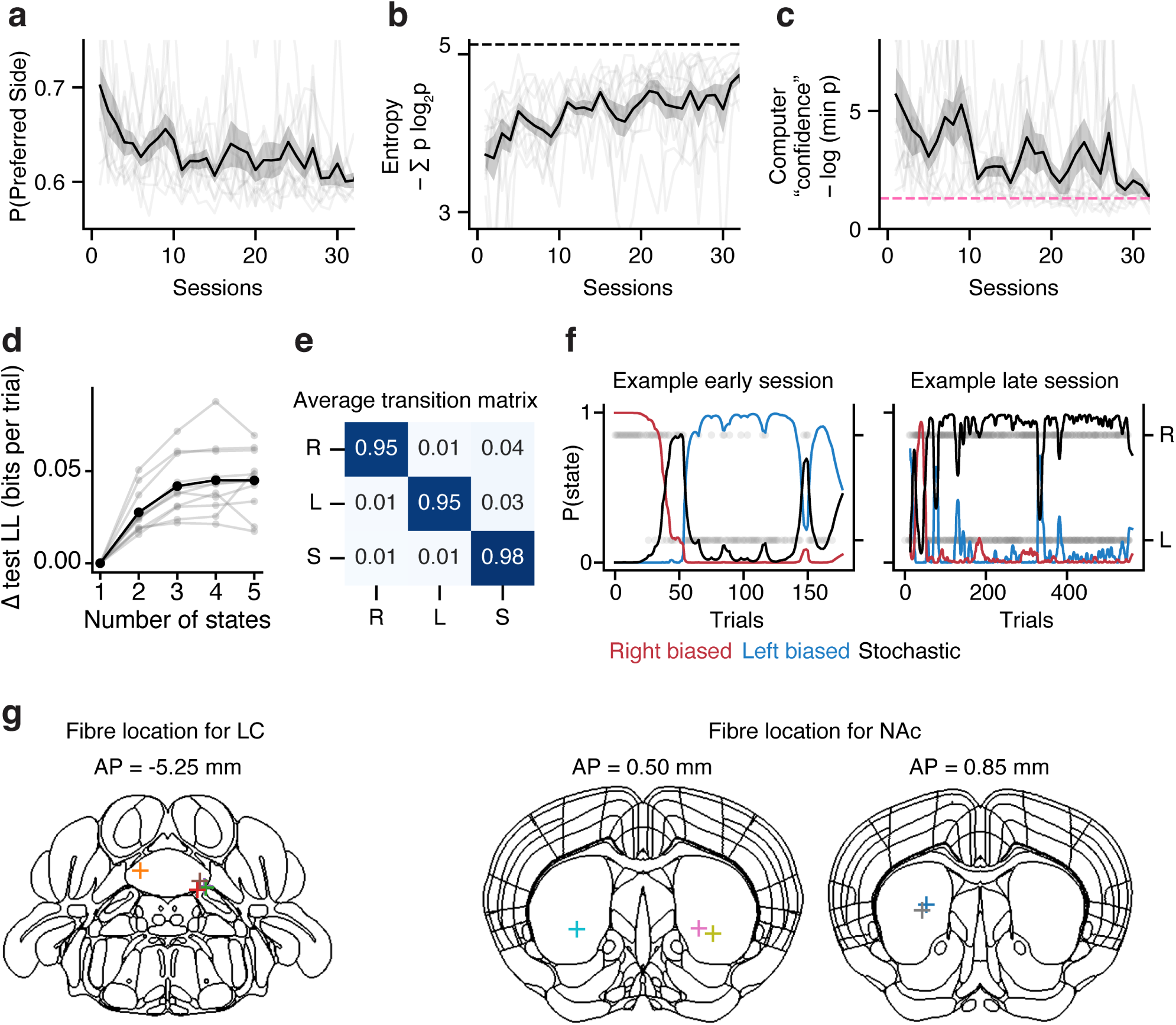
Mice learned to use more stochastic choices in the matching pennies game. **a)** Probability of choosing a biased side (left or right) across learning. **b)** 5 bit entropy across learning. **c)** Computer “confidence”, measured as the negative log of the minimum p value associated with the computer’s prediction algorithm, across learning. Pink dashed line indicates *p* = 0.05, the threshold at which the computer’s algorithm predicts left and right randomly. **d)** Difference in test log likelihood of GLM-HMM fits with different numbers of states. **e)** Average GLM-HMM transition matrix across all 11 animals. **f)** P(state) in an example early session (left) and late session (right). **g)** Locations of fibre tip for animals implanted in the LC (left) and NAc (right). Coloured crosses refer to different animals.

**Figure S2.**
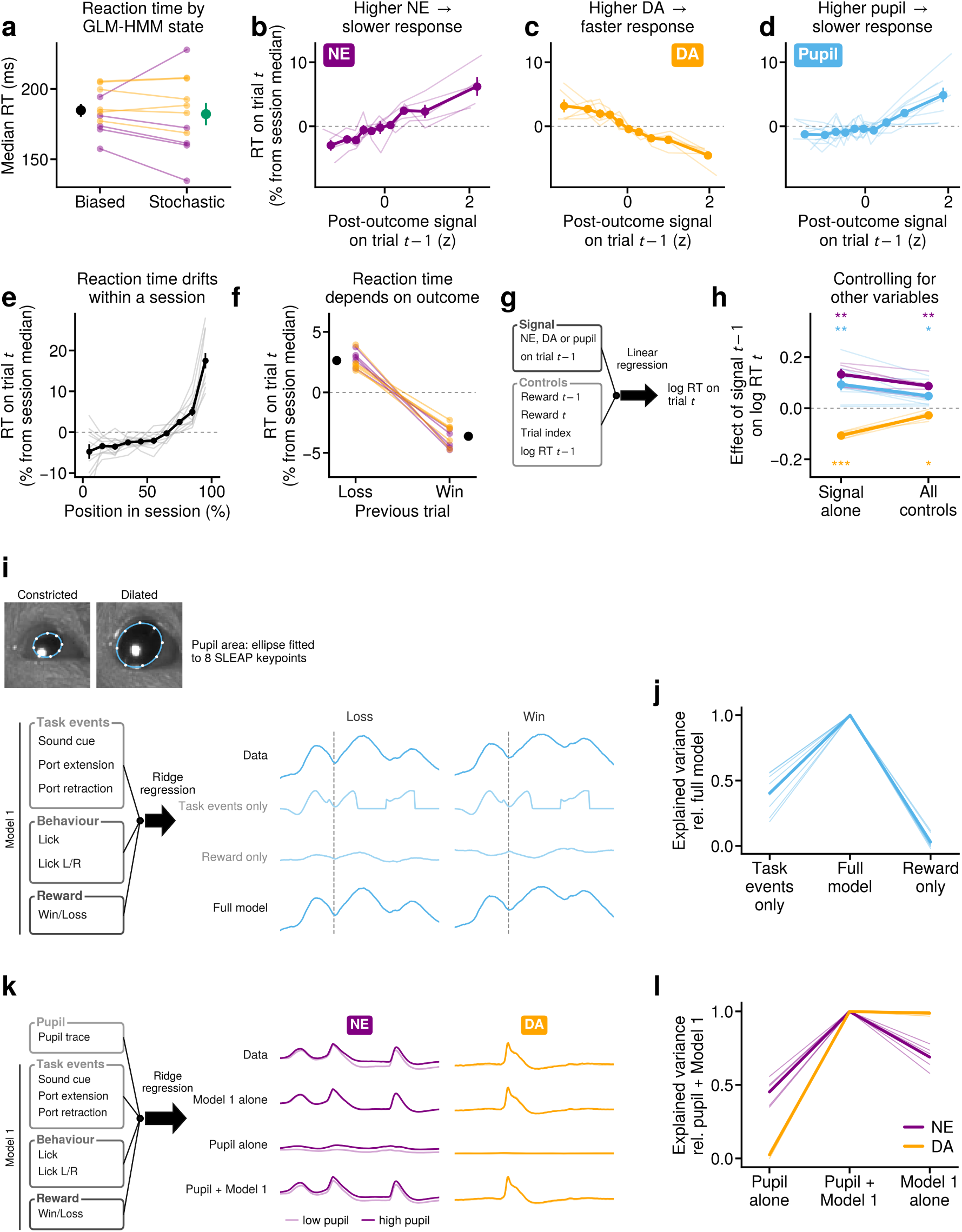
Relationship between neuromodulatory signals, arousal, and reaction time. **a)** Median reaction time across all trials in mice during the biased and stochastic states in the matching pennies task (small paired dots, n = 5 mice for dopamine and n = 5 for norepinephrine recordings). Larger dot show mean and +/-1 SEM across mice. There is no significant difference in reaction time between biased and stochastic states (p > 0.05, paired t-test). **b)** Relationship between post-outcome (2 - 4 seconds aligned to first lick) z-scored signal in norepinephrine and reaction time on the subsequent trial. Thin lines show individual mice and thick line show mean across all mice with the +/1 SEM across mice. **c)** Same as b) but for dopamine. **d)** Same as b) but for pupil response. **e)** Relationship between reaction time (percentage difference from each session’s median) and the trial index as a percentage of the total number of trials in each session. Thin lines show individual mice and thick lines show mean across all mice and +/- 1 SEM across mice (n = 11 mice). **f)** Reaction time of the current trial conditioned on whether the previous trial was unrewarded or rewarded. Lines and connected dots show individual mice (purple lines and dots show norepinephrine recordings, n = 5, orange lines and dots show dopamine recordings, n = 5). Black dots show mean across mice and line on dots show +/- 1 SEM across mice. There is a significant difference in reaction time between previous unrewarded and rewarded trials. (p < 0.01, paired t-test, n = 10 mice). **g)** Schematic of a linear regression model that models reaction time on the current trial as a function of neuromodulatory activity and other behavioural variables. **h)** Coefficient obtained from the linear regression model depicted in g) when using only dopamine, norepinephrine, or pupil post-outcome (2 - 4 seconds aligned to choice) response, or when including behavioural controls. For all three activity types, there is a significant effect of post-outcome response on the reaction time of the subsequent trial (n = 10 mice). **i)** Top: Example frames showing constricted (left) and dilated (right) pupil from the side camera in the same session. Bottom: Schematic of regression model used to predict pupil response. Traces show mean activity across mice, and the predicted response from a model using only task events, a model using only reward, and a model using all regressors. **j)** The explained variance between the predicted and observed pupil response relative to the full model in i), thin lines show individual mice and thick line show mean across all mice (n = 9 mice). **k)** Schematic of the regression model used to predict norepinephrine and dopamine activity. Traces show mean activity across mice (low pupil are obtained from trials where the mean activity of pupil is below the 33rd percentile, and high pupil obtained from trials where the mean activity is above the 66th percentile). **l)** The explained variance between predicted and observed response in dopamine (orange) and norepinephrine (purple) relative to a model with both pupil and other inputs shown in k). Thin lines show individual mice and thick line show mean across all mice (n = 4 mice for dopamine, n = 5 for norepinephrine). *** p < 0.001, ** p < 0.01, * p < 0.05.

**Figure S3.**
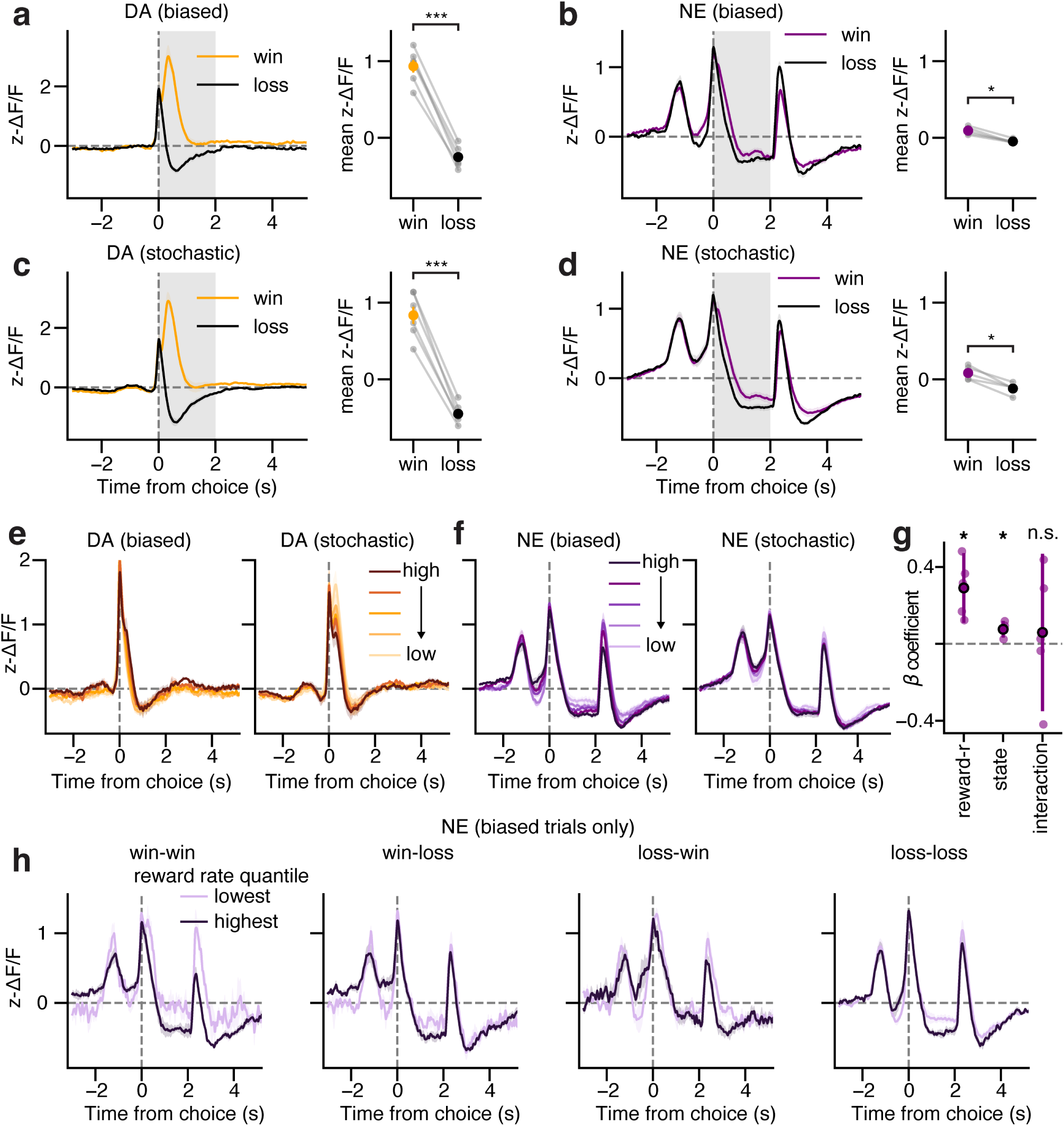
Norepinephrine reflects average reward rate when controlling for state and reward history. **a)** DA response to immediate reward outcome in biased trials. **b)** NE response to immediate reward outcome in biased trials. **c)** DA response to immediate reward outcome in stochastic trials. **d)** NE response to immediate reward outcome in stochastic trials. **e)** DA responses separated by average reward rate quantiles in the biased (left) and stochastic (right) state. **f)** NE responses separated by average reward rate quantiles in the biased (left) and stochastic (right) state. **g)** Beta coefficients from a linear regression model predicting pre-choice NE from reward_rate, state, and reward_rate:state interaction. **h)** NE responses in biased trials separated by previous outcome and current outcome for the lowest and highest reward rate quantiles. *** p < 0.001, * p < 0.05, n.s. not significant.

**Figure S4.**
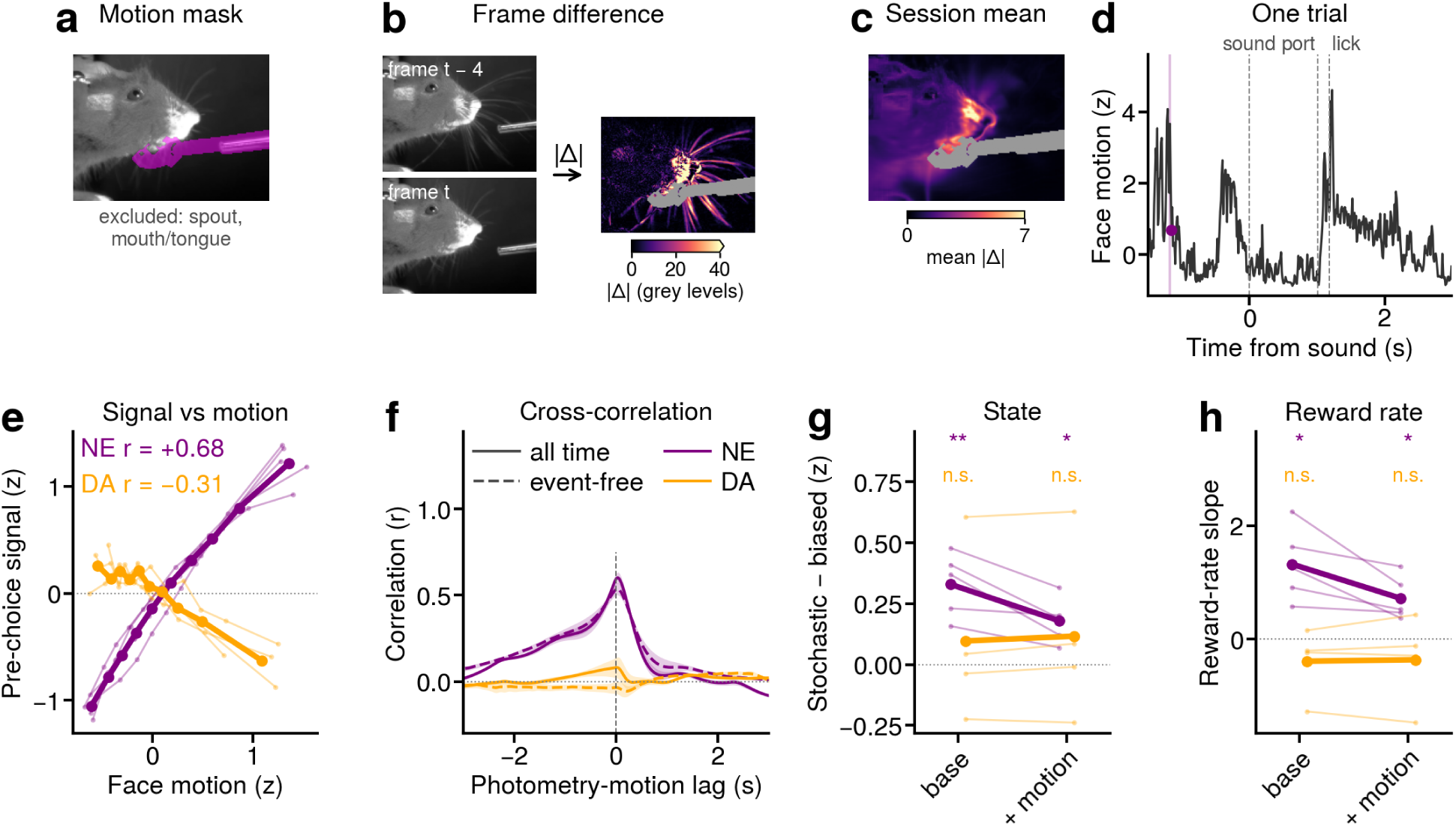
Norepinephrine representation for GLM-HMM state and average reward rate while accounting for preparatory movement. **a)** Example motion mask used to compute motion energy, with the spout and mouth area excluded from calculation to avoid introducing spout movement to the signal. **b)** Schematic showing how motion energy is quantified as the difference in absolute difference of grey level of consecutive frames. **c)** Example mean motion energy map indicating most of the motion energy is focused on the snout and whiskerpad. **d)** Example single-trial z-scored motion energy trace aligned to sound onset. **e)** Correlation between z-scored pre-choice neuromodulatory signals and z-scored face motion. Dots represent motion deciles. r values indicate the trial-by-trial Pearson correlation, averaged across mice. Thin lines represent individual mice and thick lines represent mean across mice (N = 5 mice for NE, N = 4 mice for DA). **f)** Cross-correlation between neuromodulatory signals and motion energy, during all time periods (filled line) and during inter-trial interval, where there are not task events such as sound onset or port movement (dashed line). Negative lags indicate neuromodulatory signal leading motion. Shading represents standard error of the mean across mice. **g)** The effect of state (stochastic versus biased) on pre-choice neuromodulatory signal (-2 to 0 seconds aligned to choice, z-scored) without any controls (base), and accounting for the trial-by-trial effects of facial movement (+motion). Slopes are obtained via ordinary least-square regression, and tested for significance using a one sample t-test across mice. **h)** Same as **g**, but testing the effect of reward rate. * p < 0.05, ** p < 0.01, n.s. not significant.

**Figure S5.**
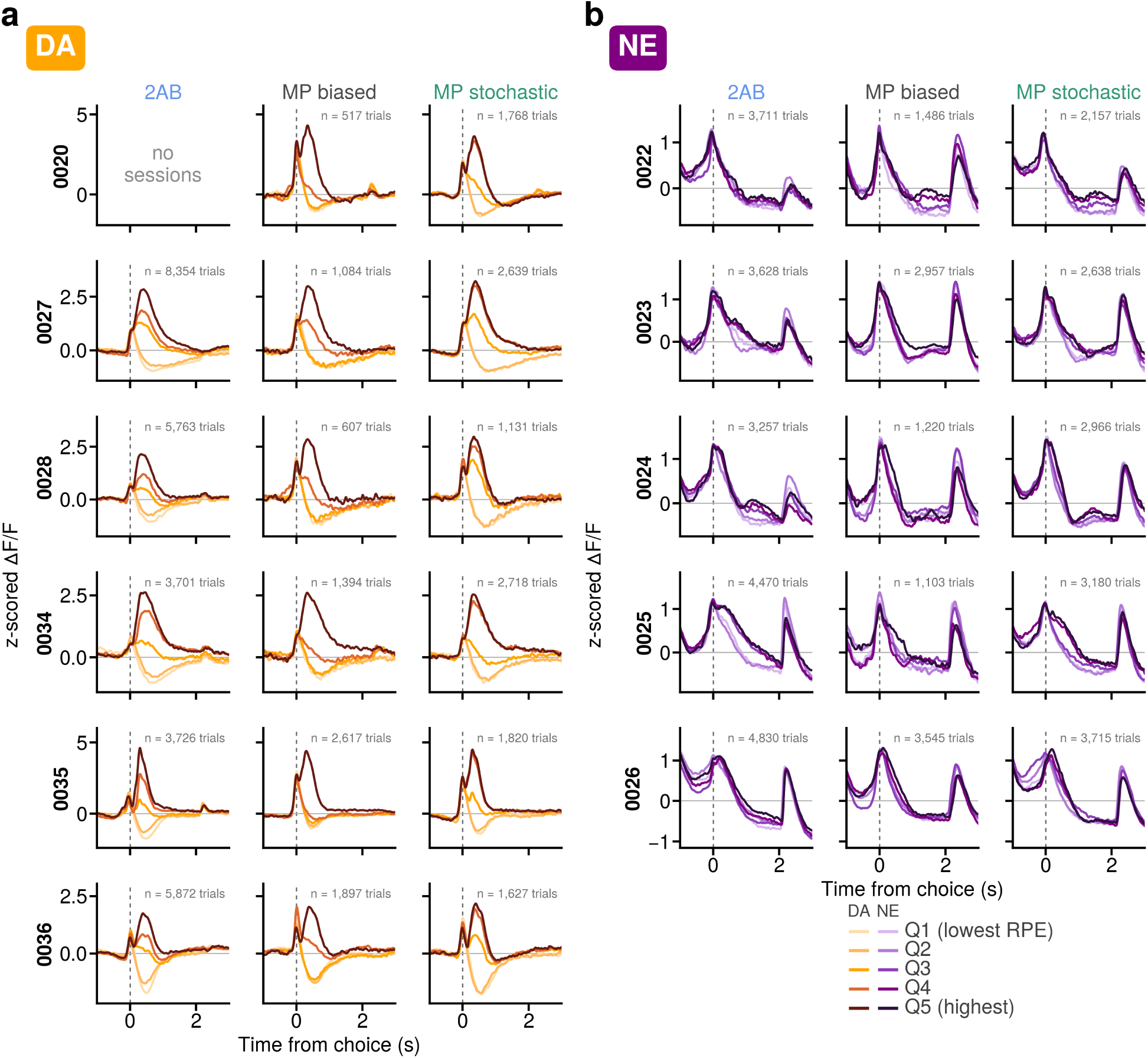
Strategy-related representations in norepinephrine when controlling for effects over learning and within-session state occupancy. **a)** Schematic where each rectangle represent a trial on a day (row), either in the biased (black) or stochastic (green) state. The left panel depicts the pooling of trials across all sessions to calculate differences in neuromodulatory response across states. The right panel depicts the calculation of difference across states first within each day, then averaging across days. **b)** NE response aligned to choice during biased (black) and stochastic (green) states, computed using trials pooled across sessions (dotted lines) or by first computing differences within session and averaging across sessions (solid lines). **c)** Effect of state difference on pre-choice (-2 to 0 seconds aligned to choice) NE (purple) and DA (orange) signal computed using pooled or within-session method. Slope is obtained via per mouse ordinary least square regression and one-sample t-test for significance of coefficient. Small dots represent individual mice and large dot represent mean across all mice (N = 5 for NE, N = 6 for DA). Error bar indicate standard error of the mean across mice. **d)** Mean trace of switch versus stay response during rewarded trials during the biased and stochastic state. Shading represent standard error of the mean (N = 5 mice). **e)** Difference in switch versus stay z-scored activity post-choice (0 – 4 seconds) during win (purple) and loss (black) trials in biased and stochastic states, either calculated using the pooling method (left) or using the within-session comparison method (right). Paired t-test across mice (N = 5). **f)** Same as e but for dopamine (N = 6 mice). **g)** Proportion of trials in the biased (black) or stochastic (green) state as a function of fraction of trials within a session, error bars show SEM across mice (N = 11 mice). **h)** Same as **g** but for reaction time during each state. **i)** Same as **g** but for proportion of miss trials. **j)** Same as **g** but for licks per trial. **k)** The proportion of trials in the stochastic state and biased state as a function of binned position within a session before (top) and after (bottom) matching for the number of trials that contribute to analyses. Lines show mean across mice (N = 11). **l)** Effect of state on the pre-choice (-2 to 0 seconds aligned to choice) neuromodulatory signal (NE = purple, DA = orange) before (left) and after (right) matching for position of trials within sessions. Small dots indicate individual mice and large dot indicate mean across mice. Error bars represent standard error of the mean. **m)** Same as **e** but comparing before (left) and after (right) matching for position of trials within sessions. * p < 0.05, ** p < 0.01, n.s. not significant.

**Figure S6.**
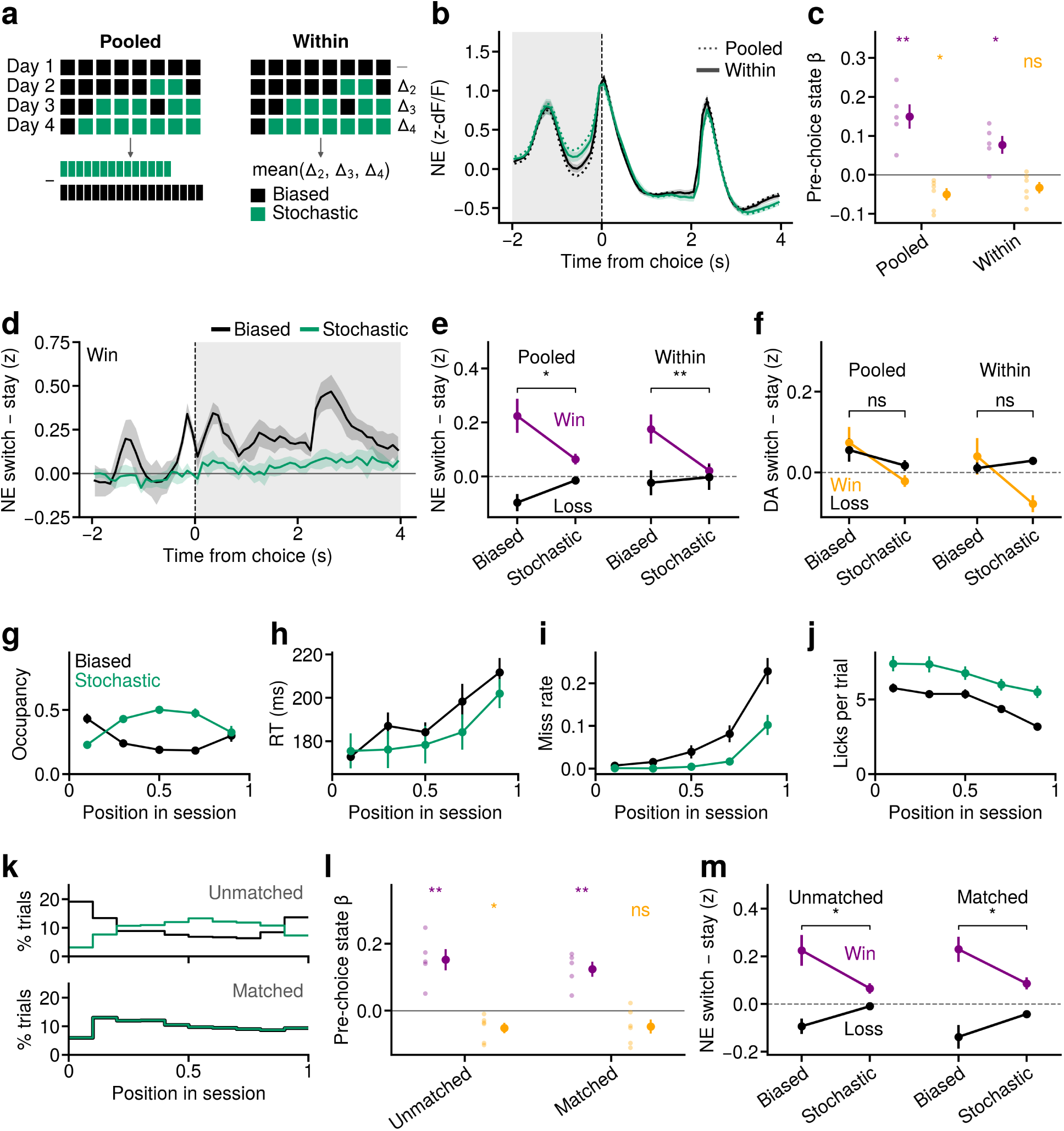
Dopamine and norepinephrine response binned by reward prediction error in each mouse. **a)** For each mouse (row), the choice-aligned dopamine response grouped by reward prediction error (RPE) quantiles during the two-armed bandit (2AB) task, and during the matching pennies (MP) task in the biased and stochastic states. **b)** Same as a) but for norepinephrine responses.

**Figure S7.**
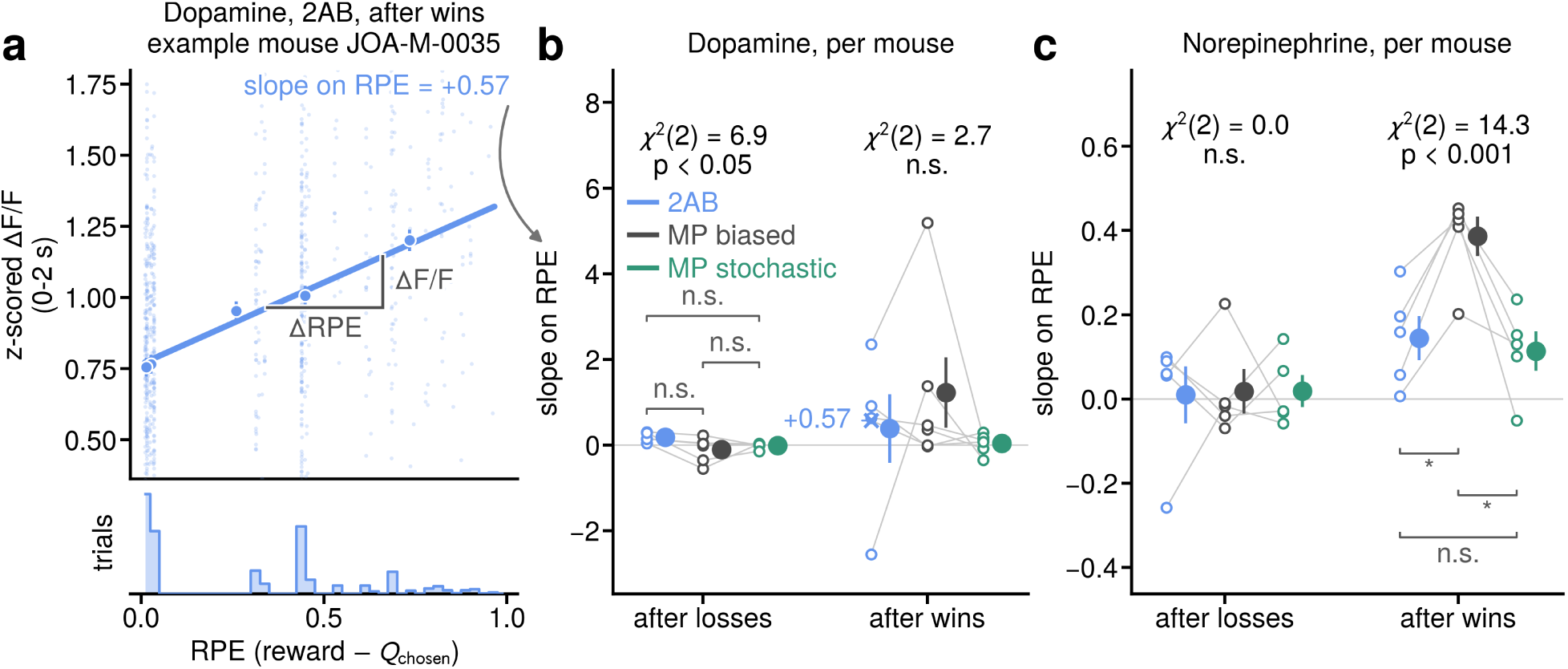
Relationship between neuromodulatory signals and RPE across tasks and states. **a)** In one example animal across all sessions of the two-armed bandit task, the fitted linear relationship between RPE and z-scored dopamine response. Small individual dots show single trials and larger dots show binned RPE and response values with equally sized bins. **b)** Fitted slope between dopamine and RPE in unrewarded (left) and rewarded (right) trials in the two-armed bandit (blue) task, and the biased (black) and stochastic (green) state of the matching pennies task. Open circles show individual mice and filled circle show mean across all mice, with line from the filled circle indicating +/- 1 SEM across mice. Cross symbol denotes the example shown in a). **c)** Same as b) but for norepinephrine.

**Figure S8.**
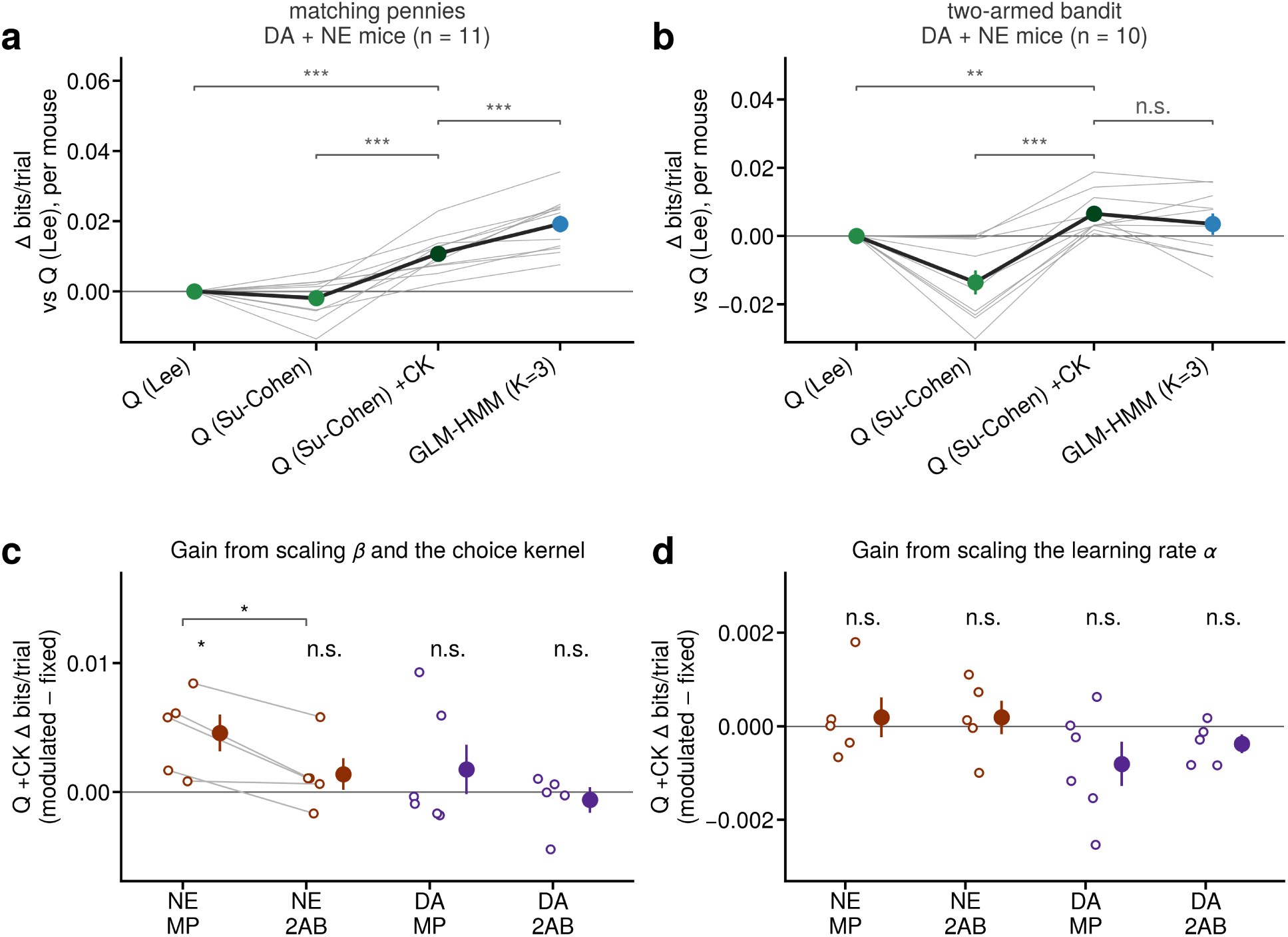
Model comparison of Q-learning + Choice Kernel model with other models. **a)** Cross-validated log-likelihood difference between Q-learning models compared to the Q-learning model in Lee et al., 2004, fit to matching pennies behaviour. **b)** Same as a) but fit to choices in the two-armed bandit task. **c)** Cross-validated log-likelihood difference between Q-learning models with choice kernel fit with neuromodulator-scaled (using pre-choice activity, -1 to 0 seconds relative to choice) temperature and choice kernel term and models with a constant temperature and choice kernel term. There is a significant increase in model performance when the temperature and choice terms are scaled using norepinephrine activity during the matching pennies task (p < 0.05, paired t-test, n = 5 mice), but not for dopamine activity and the two-armed bandit task. **d)** Same as c), but instead using the post-choice activity (2 – 4 seconds relative to choice) to scale the learning rate term.

